# Quantitative and Kinetic Proteomics Reveal ApoE Isoform-dependent Proteostasis Adaptations in Mouse Brain

**DOI:** 10.1101/2024.08.13.607719

**Authors:** Nathan R. Zuniga, Noah E. Earls, Ariel E. A. Denos, Jared M. Elison, Benjamin S. Jones, Ethan G. Smith, Noah G. Moran, Katie L. Brown, Gerome M. Romero, Chad D. Hyer, Kimberly B. Wagstaff, Haifa M. Almughamsi, Mark K. Transtrum, John C. Price

## Abstract

Apolipoprotein E (ApoE) polymorphisms modify the risk of neurodegenerative disease with the ApoE4 isoform increasing and ApoE2 isoform decreasing risk relative to the ‘wild-type control’ ApoE3 isoform. To elucidate how ApoE isoforms alter the proteome, we measured relative protein abundance and turnover in transgenic mice expressing a human ApoE gene (isoform 2, 3, or 4). This data provides insight into how ApoE isoforms affect the *in vivo* synthesis and degradation of a wide variety of proteins. We identified 4849 proteins and tested for ApoE isoform-dependent changes in the homeostatic regulation of ∼2700 ontologies. In the brain, we found that ApoE4 and ApoE2 both lead to modified regulation of mitochondrial membrane proteins relative to the wild-type control ApoE3. In ApoE4 mice, this regulation is not cohesive suggesting that aerobic respiration is impacted by proteasomal and autophagic dysregulation. ApoE2 mice exhibited a matching change in mitochondrial matrix proteins and the membrane which suggests coordinated maintenance of the entire organelle. In the liver, we did not observe these changes suggesting that the ApoE-effect on proteostasis is amplified in the brain relative to other tissues. Our findings underscore the utility of combining protein abundance and turnover rates to decipher proteome regulatory mechanisms and their potential role in biology.

For Table of Contents Only

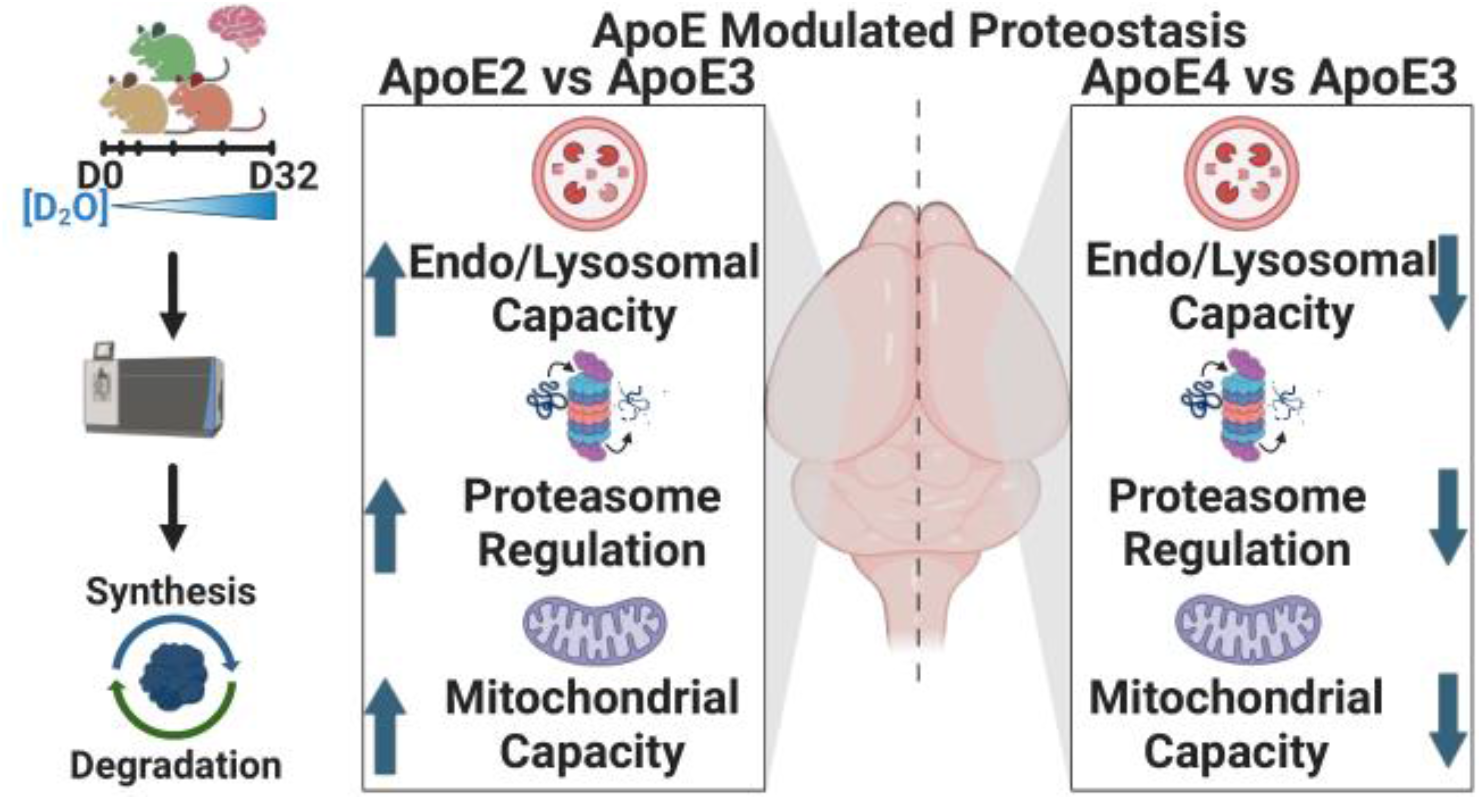

## INTRODUCTION

Apolipoprotein E (ApoE) is one of the lipoproteins used for the transport of lipids and cholesterol throughout the body. ApoE is also the primary transporter of lipids in the brain. The three major subtypes of human ApoE—ApoE2, ApoE3, and ApoE4— differ by 2 amino acids and exhibit allelic frequencies of 8.4%, 77.9%, and 13.7%, respectively. ^2, 3^ The ApoE3 allele is considered the normal or wild-type, and the behavior of the E2 or E4 isoforms differs from E3 in measurable ways. The ApoE2 protein isoform, characterized by an R158C substitution relative to the ApoE3, has been associated with decreased affinity for the LDL receptor^4, 5^, while the ApoE4 protein isoform, which features a C112R substitution relative to ApoE3, favors binding to very-low-density lipoprotein receptors^4, 5^. Thus, these seemingly minor genotypic changes may lead to profound biochemical consequences.

Both ApoE2 and E4 modulate disease risk relative to ApoE3. Ferrer et al. observed a 3 – 15-fold increase in Alzheimer’s Disease (AD) prevalence in carriers of the ApoE4 allele relative to ApoE3 carriers and a decreased risk in individuals expressing the ApoE2 allele.^6^ Although ApoE2 expression protects against AD, its expression is associated with the increased incidence of familial type III hyperlipoproteinemia—a disorder characterized by an inability to metabolize lipids including cholesterol and triglycerides.^7^ ApoE isoforms have also been implicated in the development of Parkinson’s disease^8^, vascular pathology^9^, and most recently, COVID-19 prognosis^10^.

Some mechanistic details have been identified for how the ApoE alleles modulate an individual’s risk for disease. ApoE is a transporter of amyloidβ, a widely recognized biomarker in AD development.^11^ ApoE-isoforms modulate brain mRNA expression, presumably in response to changes in lipid availability^11^ as well as direct transcriptional effects.^12^ Here we used both quantitative and kinetic proteomics to explore the impact of human ApoE genotypes in the proteome of mice. Both approaches leverage liquid chromatography and mass spectrometry (LC-MS) to identify and quantify thousands of proteins (Figure 1A).^1^ We apply a simplified kinetic model of proteostasis (Figure 1B), which combines turnover rate and concentration measurements to reveal ApoE isoform-dependent effects on protein synthesis and degradation. Our analysis identifies key brain-specific proteostasis changes, as evidenced by pathway-level changes in synthesis and degradation. Building upon a significant body of literature and this proteome scale study, we propose a unifying mechanism wherein ApoE alleles systemically impact cellular proteostasis through alterations in endosomal trafficking, mitochondrial function, and proteo-lysosomal activity.

**Figure 1.**
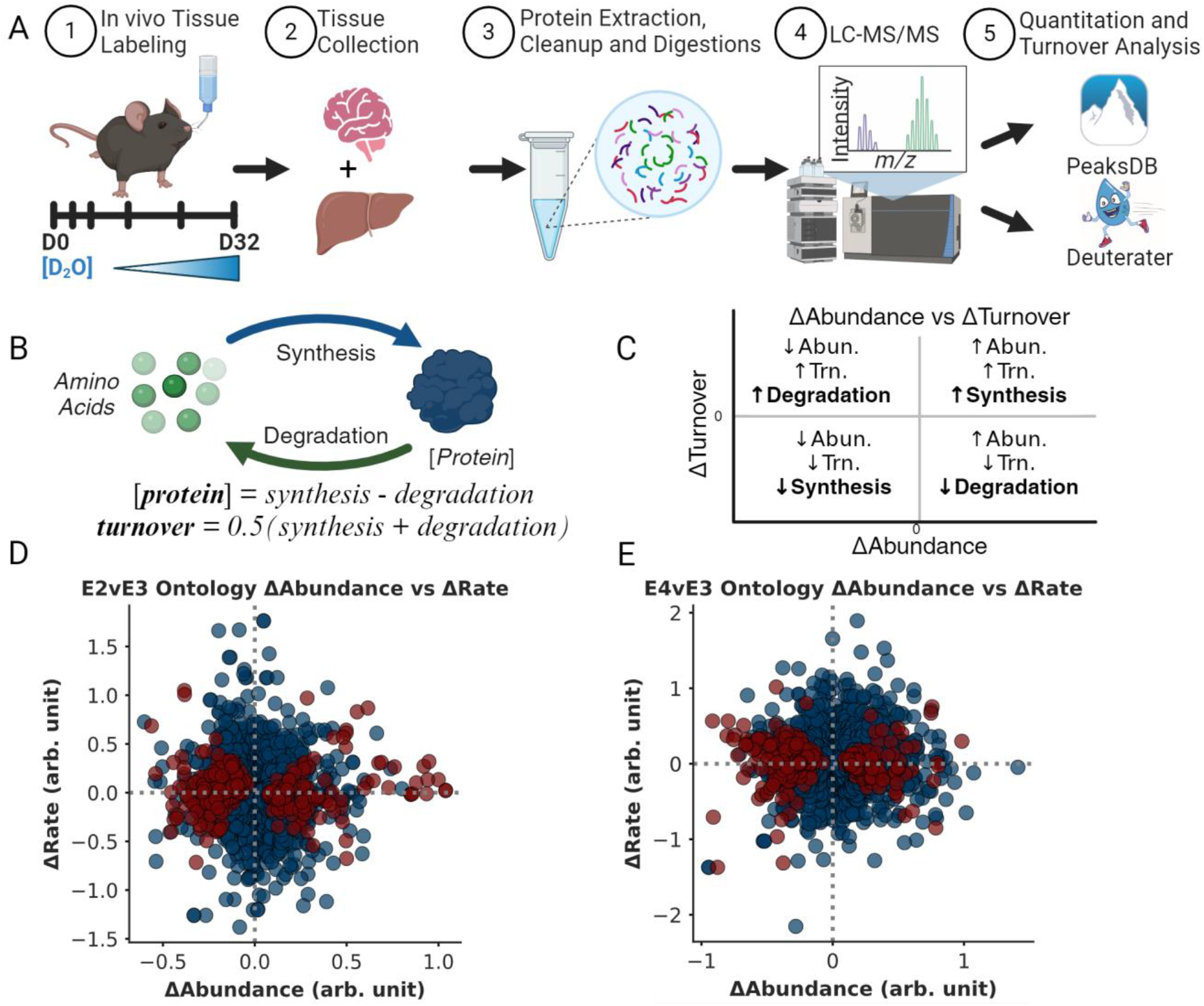
Testing for changes in regulation of the brain proteome **A**. Homozygous ApoE transgenic mice (ApoE2, E3, or E4, n = 24 each) were given 8% D2O drinking water and sacrificed at specific timepoints ranging from day 0 to day 32. Tissues were prepared using the S-Trap protocol and analyzed via LC-MS. Data-dependent acquisition was used to collect data for LFQ analysis and MS1 level data was used to calculate turnover rate values for each protein. Resulting spectra from MS/MS acquisitions were analyzed by Peaks Studio (Bioinformatics Solutions Inc.) peptide-protein identification (IDs) and quantitation while Deuterater^1^ software was used for turnover rate calculation. **B**. Proteostasis model where protein expression levels and turnover rates are a function of synthesis and degradation. **C**. Regulation of synthesis and degradation can be inferred from Δabundance (x-axis) and Δturnover (y-axis) and visualized using a proteostasis plot. **D & E**. Proteostasis plot showing 276

## EXPERIMENTAL PROCEDURES

### Experimental Design and Statistical Rationale

#### Cohort Grouping and Analysis Rationale

A total of 72 homozygous ApoE transgenic mice, with an equal distribution of female and male individuals were included. This cohort included 24 ApoE2, 24 ApoE3, and 24 ApoE4 (refer to Table S1 for details). The sample groups for protein turnover rate measurements of each ApoE genotype and gender, were two independent blocks of six mice. These six mice were selected based on the metabolic labeling duration, namely Day 0, Hour 6, Day 1, Day 4, Day 16, and Day 32 post-exposure to deuterium.

The kinetic analysis utilized peptide identifications from LC-MS/MS acquisition files to extract isotope envelope information from LC-MS (MS1 only) data. Notably, this process heavily relies on peptide retention time. To facilitate this, MS/MS data and MS data were collected within the same sample worklist. The initial four timepoints (Day 0, Hour 6, Day 1, and Day 4) were used to generate LC-MS/MS fragmentation spectra and identify peptide sequences with observed charge and retention time.

To streamline sample processing and turnover rate measurements, mice were organized into four gender-specific groups of 18 mice (n=6 per genotype, Figure S1). This grouping strategy accommodated instrument availability and minimized retention time deviations associated with extensive sample worklists. Additionally, from each group, a subset of four mice per genotype, comprising the first four timepoints (Day 0, Hour 6, Day 1, Day 4), were selected for LFQ proteomics. This selection yielded a total of 16 mice per ApoE genotype for an area under the LC curve (Abundance) fold change (FC) calculations, with an equal distribution of 8 females and 8 males in the four gender-specific groups. Thus, out of the initial 72 mice, 48 were used to generate the “Abundance” FC values (Figure S1).

To broaden proteome coverage, each brain homogenate sample was fractionated into cytosolic and membrane components, which were prepared and analyzed separately using the workflow described below. This fractionation led to the creation of eight datasets for our analysis. Each dataset underwent individual processing using the Peaks Studio software (Bioinformics Solutions Inc.) for protein abundance and Deuterater software ^1^ for turnover rate measurements. Protein-level abundance fold change relative to control (FC values), turnover rate FC values, and statistical analysis (P-value) for each comparison (e.g., E2vsE3) were calculated for each dataset independently to minimize inter-set variance caused by sample prep discrepancies, instrument noise, buffer compositions, and sample run variables. It is worth noting that due to problems in sample processing, the Hour 6 sample was omitted from a single ApoE4 dataset (D16 was substituted for LFQ analysis), and Day 4 was omitted from a single ApoE3 LFQ dataset (See Table S1, Figure S1).

While FC and P-value calculations were conducted at the protein level, this study mainly focuses on how proteins with shared functional characteristics are regulated in an ApoE isoform-specific manner. To achieve this, the StringDB multiprotein tool^13^ was employed to identify functional groups (ontologies) represented in the final data sets (Abundance FC, Turnover FC). Every protein Abundance FC value was calculated with a minimum of three biological abundance measurements in experimental (ApoE2, or ApoE4) and control (ApoE3); see the ‘Protein ΔAbundance Analysis’ section for more details. The null hypothesis (H0) posited that proteins’ collective gene expression ratio in an ontology would remain unchanged (H0: Abundance FC = 1) across ApoE genotypes. Consequently, we tested the alternative hypothesis that ApoE genotype alters the regulation of functionally related protein groups (Ha: Abundance FC ≠ 1) using a one-sample t-test. This analytical approach captured changes occurring across the broader functional proteome rather than focusing solely on identifying individually significant proteins. Python code created for both protein- and ontology-level calculations is available in the GitHub repository, as detailed in the Supplementary Data section of this paper.

#### Proteostasis Model and Analysis Rational

A protein homeostasis model must account for common sources and sinks of protein mass (Figure S2). In this model we assume there is a large circulating pool of free amino acids affected by diet, metabolites, and waste expulsion. Amino acids become the precursors for protein synthesis in an initial tRNA charging step, which then polymerize in an mRNA-dependent step before folding into functional proteins. Multiple competing processes subsequently influence the resulting protein concentration. First, degradation returns the protein to the constituent amino acids in the free pool. The functional proteins may also transition (reversibly) into an aggregate/condensate state that undergoes a separate degradation process. Finally, protein concentration may be affected by importing or exporting proteins.

Our goal is to use chemical kinetics and translate this diagram into a mathematical model with a few tunable parameters identifiable from experimental data. Unfortunately, a complete mathematical translation of this system leads to a model with too many parameters to draw meaningful conclusions. We, therefore, make several simplifying assumptions regarding which processes are dominant to restrict the parameters to an identifiable subset.

First, the mice in this study are healthy adults, so we assume the protein concentrations are in steady-state with no protein aggregate. We assume that the pool of free amino acids is large so that the rate bottleneck in the tRNA-charging/synthesis steps is synthesis and that import/export is negligible. We also assume the protein pool is well mixed, assuring the random selection of protein for degradation (unregulated). Although a reasonable starting point for the present study, we acknowledge that these assumptions are poor for surprisingly large sections of the proteome where reversible aggregation ^14, 15^, multistage regulation of synthesis rates ^16-18^, exchange of protein subunits in complexes ^19, 20^ and nonrandom degradation ^21-24^ are biologically important. There has yet to be presented a standardized model that successfully accounts for all of these confounding variables, so we elect to maintain the aforementioned assumptions as a starting point for the modeling. Previous literature reports have presented mathematical modeling of protein turnover rates using similar starting points.^25-32^ This then results in Equation 1 where the time-dependent change in a protein concentration is the difference between the synthesis and degradation rates.

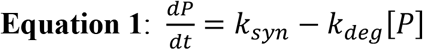

This model assumes that the concentration of an individual protein ([P]) in every location is under the control of a zero-order synthesis rate (k_syn_) and a concentration-dependent degradation (k_deg_) step. The assumption of zero-order synthesis suggests that the precursor is stable and unresponsive to protein concentration, while a first-order rate for degradation suggests that there is no regulation of degradation other than protein concentration. In general, the rates k_syn_ and k_deg_ need not be constant as they are under the control of numerous exogenous factors that may vary in time. However, we now formalize our final assumption: protein homeostasis. This assumption is that the multiple processes regulating each protein concentration are in dynamic equilibrium so that these rates are constant for a given experimental condition. Under these assumptions, both synthesis and degradation for a given protein are equal, ensuring that the number of proteins produced is equal to the number of proteins lost. Therefore, during the measurements d[P]/dt=0, leading to the relationship:

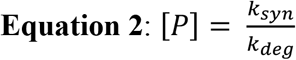

Our hypothesis is that the expression of the ApoE polymorphysms (ApoEx = ApoE2 or ApoE4) creates a unique steady state or proteostasis across the proteome that can differ from the concentration of the human wild-type control (ApoE3). The change in protein abundance (Equation 3) between the two conditions allows us to infer how the ratio of the rates is changed in the experimental cohorts, but neither parameter is individually identifiable.

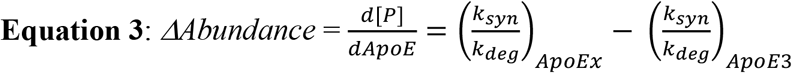

Using metabolic isotope labeling (Figure S2) we can add rate information that will distinguish between changes in synthesis and changes in degradation. Assume that at t=0 our model simplifications are true, but that the amino acid pool is replaced with a deuterated version. Proteins synthesized after t=0 are isotopically labeled, and we can measure the time-dependent replacement of old unlabeled for new labeled proteins. To make this mathematically explicit, we denote the concentration of normal proteins by [P] and the concentration of deuterated proteins by [P^D^]. These two concentrations now satisfy the initial value problems.

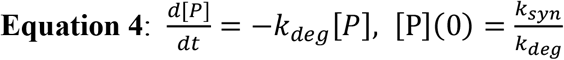

and

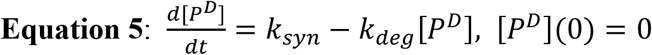

These are true because normal proteins are no longer being synthesized (Equation 4) and [P^D^] have no initial concentration (Equation 5). These ordinary differential equations can be solved in closed-form using standard techniques. The solutions are:

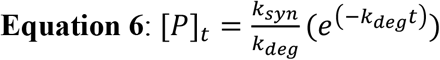

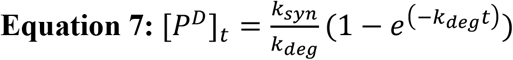

Notice that these equations satisfy [P]_t_ + [P^D^]_t_ = *k*_*syn*_*/k*_*deg*_, which is independent of time as it must be in homeostasis. However, the measurable fraction of deuterated protein over time is given by.

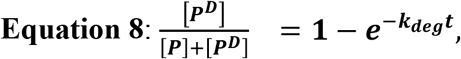

Equation 8 seems to suggest that the degradation rate is the measurable driving force behind the turnover of old protein and the replacement by labeled protein. However, because the processes of synthesis and degradation are exactly balanced in the proteostasis condition, we can just as easily identify the turnover rate as the per-molar synthesis rate: *k*_*syn*_/[*P*] or as it is commonly called fractional synthesis^33^. It is important to emphasize that these rates are only properly defined in homeostasis. Because there is no assurance that homeostasis is equally applied to all proteins simultaneously^18, 21, 34^, we find it conceptually preferable to define the turnover rate as the mean of the per-molar synthesis and degradation rates:

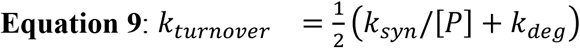

As stated above, each experimental mouse cohort will have a unique homeostasis with a protein-specific synthesis and degradation rate. Using ApoE3 as our normal control we can assess how the average of the synthesis and degradation rates have changed with the E2 and E4 polymorphisms (ApoEx).

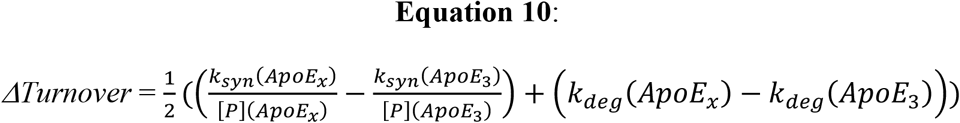

This means that if, for example, the ApoE polymorphism increases a proteins concentration (+*ΔAbundance*, Equation 3) the *ΔTurnover* (Equation 10) will highlight whether the change in proteostasis was driven by an increase in *k*_*syn*_*/[P](ApoEx)* or a decrease in *k*_*deg*_*(ApoEx)* because the sign of *ΔTurnover* will be different for each possibility. Together the unique *ΔAbundance* and *ΔTurnover* for each protein identifies whether the differences in proteostasis are primarily due to changes in synthesis or degradation. Graphing these values produces a plot where each quadrant (Figure 1C) has meaning. For example, a positive x-axis (+Δabundance) and y-axis (+Δturnover) suggest that synthesis increases (Syn↑). Conversely, a protein with lower expression levels (-Δabundance) between ApoE genotypes, could result from less synthesis (Syn↓) if turnover rate decreases (-Δturnover) or more degradation (Deg↑) if the protein turnover rate increases (+Δturnover). Since each measurement has independent noise, a nonrandom grouping of multiple proteins from a functionally related ontology within a quadrant is an important metric of confidence that the cell is regulating protein expression to change biochemical functions (Figure 1D and 1E).

#### Mouse Handling

All animal handling experiments were authorized by the Brigham Young University Institutional Animal Care and Use Committee (IACUC protocol #191102). The mouse model employed for this study consists of C57BL/6 transgenic mice with homozygous genotypes for each of the three human ApoE alleles (ApoE2, ApoE3, ApoE4, n=24/allele, see Supplemental Table S1) under the GFAP promoter (JAX# 004632, 004633, 004631). Notably, this model has provided valuable insight into genotype-specific effects of ApoE in a large number of other experiments^35-46^. This study does not encompass differences from wild-type mice. The transgenic mice were selected with deliberate focus on ApoE isoforms rather than wild-type conditions, or age differences. The findings reported in this publication use fold change relative to ApoE3 to minimize the GFAP promoter variable as reported previously^47-49^. Because mouse ApoE has a low sequence identity (77%) and a different transcription promoter, the human ApoE3 model is the best control for comparison. While we recognize the limits of a transgenic model, this study provides valuable identification of *in vivo* patterns which can be confirmed in future ApoE knock-in mice models and human studies. These results refer solely to the effects of ApoE isoform differences, rather than Alzheimer’s Disease. Any claims regarding Alzheimer’s disease are made solely to highlight similarities between current ApoE/AD research and our observations to create a holistic mechanistic hypothesis.

Mice were randomly selected for replicate designation and timepoint based on availability. They were all 6–8-month-old, retired breeders with no signs of disease or neurological dysfunction. There were no exclusions among this group. Specific cohort denominations and animal numbers can be found in Table S1. Blinding was not used during any portion of this experiment as it was necessary to compare groups at each point. Mice were housed together in the same room of the facility at the same time. Mice had *ad libitum* access to water and standard nutritional rodent feed (Teklad 8604) while housed in a temperature-controlled environment of ∼24 ^°^C. This environment included a 12-hr circadian cycle. To initiate turnover rate measurements, mice received an intraperitoneal (IP) injection of sterile D_2_O 0.9% w/v saline (35 μl/g body weight) calculated to increase internal D_2_O concentrations to an initial 5% of overall water weight (w/w). Mice were then given 8% D_2_O as the sole hydration source for the remainder of the experiment. This was done to maintain overall internal water at 5% D_2_O enrichment. Mice were sacrificed according to the following timepoints post IP injection: day 0 (no D_2_O injection), hour 6, day 1, day 4, day 16, and day 32. Mice were euthanized via CO_2_ asphyxiation followed by bilateral thoracotomy. Blood was collected via cardiac puncture for D_2_O enrichment calculations. Brains were divided sagittally into respective hemispheres. Relevant organs including brain and liver were flash frozen on blocks of solid CO_2_. Tissues were stored frozen at -80 °C until processing.

#### Tissue Preparation

Singular brain hemispheres and liver sections were homogenized in lysis buffer (25mM Ammonium Bicarbonate treated with diethylpyrocarbonate and ThermoScientific Halt Protease & Phosphatase Inhibitor Cocktail) for 60 sec using a MP FastPrep-24 homogenizer. Homogenized samples were centrifuged for 15 minutes at 14,000xg to separate them into cytosolic and membrane isolates. The membrane pellet was resuspended in lysis buffer and centrifuged for 15 minutes at 14,000xg a total of three times to remove cytosolic components. Each fraction was resuspended in 5% SDS. Aliquot concentration was measured via a Pierce™ BCA Protein Assay Kit purchased from ThermoFisher Scientific, and 50 μg of protein were prepared according to S-Trap^™^ documentation (cytosol and membrane fractions were prepared separately). Proteins were digested with trypsin Lys/C overnight at 36°C. Resultant peptides were dehydrated in a ThermoScientific Savant SPD131DDA SpeedVac Concentrator and resuspended at a final concentration of 1 μg/μL in buffer A (3% acetonitrile, 0.1% formic acid).

#### LC-MS

Samples were separated and measured via liquid chromatography-mass spectrometry (LC-MS) on an Ultimate 3000 RSLC in connection with a Thermo Easy-spray source and an Orbitrap Fusion Lumos. Peptides were pre-concentrated with buffer A (3% acetonitrile, 0.1% formic acid) onto a PepMap Neo Trap Cartridge (particle size 5 μm, inner diameter 300 μm, length 5 mm) and separated with an EASY-Spray™ HPLC Column (particle size 2 μm, inner diameter 75 μm, length 25 mm) with increasing buffer B (80% acetonitrile, 0.1% formic acid) gradient:

0-5 min, 0 to 5% B; 5-87 min, 5 to 22% B; 87-102 min, 22 to 32% B; 102-112 min, 32 to 95% B; 112-122 min, 95% B; 122-125 min, 95 to 2% B; 125 to 127 min, 2% B; 127-129 min, 2 to 100% B; 129-132 min, 100% B; 132-133 min, 100 to 2% B; 133-135 min, 2% B; 135-137 min, 2 to 100% B; 137-140 min, 100% B; 140-142 min, 100 to 0% B; 142-144 min, 0% B.

The MS-based data-dependent acquisition method was set to a 3 second cycle time. MS1 scans were acquired by the Orbitrap at a resolution of 120,000. Precursors with a charge > 1 and < 6 were selected for MS2 fragmentation. MS2 scans of CID precursor fragments were detected with the linear ion trap at a scan rate of 33.333 Da/sec with a dynamic injection time. CID collisions were set to 30% for 10ms. A 60 second dynamic exclusion window was enabled; isotopes and unassigned charge states were excluded. The deuterium labeling information was collected separately in an MS1-only acquisition with the Orbitrap at a resolution of 60,000 as previously described by Naylor et al.^1^ The mass spectrometry proteomics data have been deposited to the ProteomeXchange Consortium via the PRIDE partner repository with the dataset identifier PXD044460

#### Raw Data Processing for Peptide Identification and Label-free Quantitation

Raw files were searched against the 2022 Uniprot/Swissprot *mus musculus* FASTA (containing 17144 entries) using Peaks Studio v.11 (Bioinformics Solutions Inc.) for label-free quantitation (LFQ) analysis. During the data refinement step, the feature “associate feature with chimera [DDA]” was selected to deconvolute scans with co-eluted isobaric peptides. The parent mass error tolerance was set to ±15 ppm and the fragment mass error tolerance was set to 0.5 Da. Cysteine carbamidomethylation was set as a fixed modification, and both methionine oxidation and pyro-glu from glutamine were set as variable modifications in the search. Digest mode was set to semi-specific for the trypsin-lysC enzyme mix allowing for ≤ 3 missed cleavages and the peptide length range was set to 6 – 45 amino acids. The false discovery rate (FDR) for peptide matches was set to 1%, and protein ID significance was set to -10log(P-value) ≥ 15 for each identified protein.

Peaks Studio (Bioinformics Solutions Inc.) was also used to search raw files for use in Deuterater^1^ software. The raw files were searched against the 2021 Uniprot/Swissprot *mus musculus* FASTA (containing 17144 entries). Peptide searches were performed using trypsin/lysC semi-specific digest with a tolerance of ±20ppm and missed cleavages ≤ 3. Carbamidomethylation was set as a fixed modification and pyro-glu from glutamine and methionine oxidation were set as variable modifications. Within the Peaks Studio DB module, proteins were identified with two or more unique peptides at an FDR of 2% and significance was set to -10log(P-value) ≥ 15 for each identified protein.

#### Protein ΔAbundance Analysis

The group of mice used in this paper were divided into two male and two female groups for analysis. Each group produced a dataset for cytosolic proteins and another dataset for membrane proteins. Please refer to the *Experimental Design and Statistical Rationale* section for more information resulting in a total of eight datasets.

Data filtering, normalization, and quantitative calculations were performed independently for each dataset following standardized metrics for data quality and analysis following the process described by Aguilan et al. ^50^ Each Peaks Studio DB *protein*.*csv* output dataset contains the proteins identified in the analysis and the expression values (relative abundance) for each protein in each sample are labeled as “Area”. This output was filtered to retain only the top proteins in each protein group and proteins with at most one missing protein “Area” value per genotype (i.e., n – 1/genotype/dataset). Subsequently, protein “Area” values in the dataset underwent log_2_ transformation. The distribution of these protein “Area” values was mean centered by subtracting by the average protein “Area” from each protein “Area” within the sample. To ensure comparability across samples, the distribution width was also normalized between samples by calculating the correlation slope between these total average protein “Area” values across all samples and the individual sample values. Each protein “Area” in a sample was then divided by the corresponding sample slope. For samples with a missing protein “Area” value, imputation was carried out using the scikit-learn KNN imputer function module in python with the two closest neighbors.^51^

Protein fold change (FC) values, which represent the relative change in protein abundance values (“Area”) compared to a reference, were calculated, and used as a metric of change in abundance (Δabundance). For this study, FCs were calculated for protein expression values in ApoE2 mice and ApoE4 mice with ApoE3 expression values as reference, respectively. As per Aguilan et al.’s methodology, an F-test was employed to assess the variance between protein expression values before performing p-value calculations for statistical significance. To evaluate the statistical significance of expression levels in each comparison, a two-sample independent t-test (homoscedastic) was employed for proteins with an insignificant F-test result and a two-sample independent t-test (heteroscedastic) for proteins with a significant F-test result. Both the F-test and t-test calculations were conducted with the Scipy python package.^52^

Protein FC values were averaged across all datasets for each respective comparison. This produced a single set of “Area” (expression value) FCs for each comparison. Please note that both the ApoE2 vs ApoE3 (E2vsE3) comparison and the ApoE4 vs ApoE3 (E4vsE3) use the same list of quantified proteins. As outlined by Van den Berg, protein FC values from individual comparisons were range scaled using the following formula prior to ontology exploration ^53^:

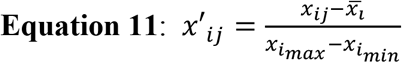

Where *x’*_*ij*_, *x*_*ij*_, 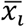, *x*_*imax*_, and *x*_*imin*_ are the scaled FC value, non-scaled FC value, mean FC, largest FC, and smallest FC, respectively. Range scaling was selected because it captures relative change in protein expression while considering the full range of values specific to the dataset. These scaled FC values will be utilized in functional analyses as described in the *Ontology-level Calculations* section below. The python script created for the steps outlined in this quantitative analysis can be found in the GitHub repository as detailed in the *supplementary data* section of this paper. The change in abundance was validated in samples using Data Independent Acquisition (DIA)^54^ to test for reproducibility of the ontology level changes (Supplementary table S6).

#### Protein ΔTurnover Rate calculation

Protein turnover rate values were calculated using Deuterater^1^ v5. This software uses an accurate mass and time database to extract peptide isotope patterns from LC-MS (MS1) centroided data utilizing feature identifications (e.g. retention time, mass, peptide ID, etc.) obtained from MS/MS data (refer to the *Raw Data Processing for Peptide Identification and Label-free Quantitation* section above).

Isotope patterns were extracted from MS1 raw data with an extraction retention time window of 1.5 min and an m/z error limit of ≤ 30 ppm. The n-value represents the number of available positions on a peptide where deuterium can replace hydrogen. In the *theory generation* step, peptides with data missing in an extracted file are removed, and the n-value is calculated for remaining extracted peptides based on known quantities for each amino acid^55, 56^. Subsequently, *Fraction New* measures the amount of turnover rate for each peptide in a file by calculating changes in neutromer abundance and spacing^1^. These calculations were performed using the average between M0 and the highest isotope peak for peptides meeting specified criteria, including peptide n-values greater than 5, a minimum peptide sequence length of 6, and a minimum allowed M0 change of 0.04. In the *Rate Calculation* stage, the data from the *Fraction New* step is fitted to a kinetic rate curve using **Equation 8** from our proteostasis model. Turnover rates were calculated for peptides that met a specified criterion, including a minimum of 3 non-zero peptide timepoints, and measurement deviation of less than 0.1, as previously reported^19^. The asymptote value is assumed to be 1 in the first iteration of analysis for proteins, but not for lipids where multiple pools of the same lipid are frequently observed^57^.

After the Deuterater^1^ analysis, all proteins with a valid turnover rate value (Rsq ≥ 0.6, combined unique peptides > 1, combined rate > 0) grouped by allele cohort, and the average turnover rate value was calculated for each protein in the cohort, respectively. These protein turnover rate values were log_2_ transformed, and the protein turnover rate FC was calculated as the difference of the log_2_ rates. The ApoE2 mice and ApoE4 mice were compared to the reference ApoE3 mice, resulting in a single set of protein turnover rate FCs for each comparison. To standardize the protein turnover rate FCs, auto scaling ^53^ was applied, where *x’*_*ij*_, *x*_*ij*_, 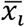, and *s*_*i*_ are the scaled turnover rate FC value, non-scaled turnover FC rate value, mean turnover FC rate and turnover rate FC standard deviation, respectively:

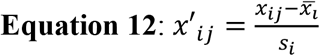

Auto scaling was implemented because it considers both the mean and standard deviation to reduce the effects of outliers and variation in the data while preserving the ability to focus on small changes. It is important to note that because of the signal to noise requirements fewer proteins had valid turnover rate FC values than quantifiable abundances. Consequently, proteins with turnover rate FCs represent a smaller subset population in comparison proteins with expression value FCs calculated from “Area” values. These protein turnover rate values were used for ontology analysis as outlined in the *Ontology-level Calculations* section below. For further reference, the python script created to process the *calculated_rates* output from Deuterater^1^ can be found in the GitHub repository, as detailed in the supplementary data section of this paper.

#### Ontology-level Calculations

The StringDB^13^ multiprotein tool was employed to calculate group FC values for functionally-related protein groups (ontologies) regardless of statistical significance. To streamline the analysis and reduce the number of redundant term ID (ontologies), ontologies were selected only from the following established: *GO Process, GO Function, GO Component, KEGG, Reactome*, and *WikiPathways*. To quantify the representation of each ontology, the “observed gene count” was divided by the “background gene count” to calculate the “Ontology_coverage (%)” for each ontology. Only ontologies with ≥25% were included in this analysis. This latter criterion ensures that the identified ontologies are sufficiently represented in the data (Table S4 and S5).

The “matching proteins in your network (labels)” was used to associate each observed protein in the ontology with the calculated “Area” FC and turnover rate FC, respectively, for both the E2vsE3 and the E4vsE3 comparison. Next, the average protein “Area” FC and turnover rate FC was calculated for each identified ontology by averaging the FC values of proteins within that category. This step summarized the collective expression and turnover rate changes of proteins within specific functional groups for each comparison.

To assess the statistical significance of the FC values within each ontology, a one-sample t-test was performed with null hypothesis (H0) stating the average ΔAbundance (**Equation 3**) for the ontology is equal to 0, and the alternative hypothesis (Ha) indicating that it is not equal to 0. This statistical test is used to determine whether the observed changes in protein expression for the ontology were statistically significant. To account for multiple comparisons and maintain a controlled false discovery rate (FDR), the Benjamini-Hochberg correction (BH-PV) was calculated for the resulting p-values (FDR = 0.25). Ontologies with a BH-PV < 0.05 were considered statistically significant. In the case of highly similar ontology with significant changes, the ontology with the most proteins was selected to represent the results. The Python code used to analyze StringDB and calculate the FCs can be found in the GitHub repository, as detailed in the supplementary data section of this paper.

## RESULTS

### Proteome Ontology Analysis

In our analysis, we identified 4,849 proteins in the brain tissue across the three ApoE-isoform groups (n = 47). We determined abundance and turnover rate FCs for comparisons for these proteins: ApoE2 vs. ApoE3 (E2vsE3) and ApoE4 vs. ApoE3 (E4vsE3). Here, ApoE3 serves as the reference ‘normal’ control. We quantified 3,532 abundance FCs for both the E2vsE3 and E4vsE3 comparisons (Supplementary Table S2). A smaller number of turnover rate FCs were obtained: 1,430 for E2vsE3 and 1,405 FCs for E4vsE3 (Supplementary Table S2) because of the more rigorous statistical filtering criteria.

For this analysis, we focused on ontologies from six databases: *GO Function, GO Component, GO Process, WikiPathways, Reactome*, and *KEGG*. Using the StringDB results, we calculated the average abundance FC (Δabundance) and turnover rate FC (Δturnover) for the proteins observed in each ontology (Please refer to the *Ontology-level Calculations* in the methods section). This yielded ∼2700 ontology-level comparisons between average Δabundance and Δturnover calculations for both E2vsE3 and E4vsE3 (Supplementary Table S4). The interpretation (Figure 1C) relies on the traditional understanding of protein turnover, contextualizing changes in protein expression. It offers a lens to assess the variances in the steady states of ApoE genotypes^58^. Using a one-sample t-test, we discerned which ontologies deviated significantly from a median Δabundance of 0. In the E2vsE3 comparison, we identified 284 protein ontologies with notable Δabundance (BH-PV < 0.05) (Figure 1D). For the E4vsE3 comparison, 287 protein ontologies had significant Δabundance (BH-PV < 0.05) (Figure 1-E).

The box plots in Figures 2 through 5 encapsulate the Δabundance and Δturnover for ontologies with marked Δabundance shifts. To maximize visibility and to accommodate for space limitations, these boxplots do not contain outlier points but supplementary Figures 2 through 5 contain the boxplots with outlier points. Given that some ontologies are repetitive, proteins depicted in the box plots might appear in multiple ontologies with analogous names/functions. When faced with such redundancies, we typically chose the ontology with superior coverage (Observed/Total) for representation. As a convention, each ontology is presented in an “*ontology name (n)*” format, where (n) indicates the count of quantified proteins within that ontology. Overlap between similar ontologies is shown in the heatmap.

**Figure 2.**
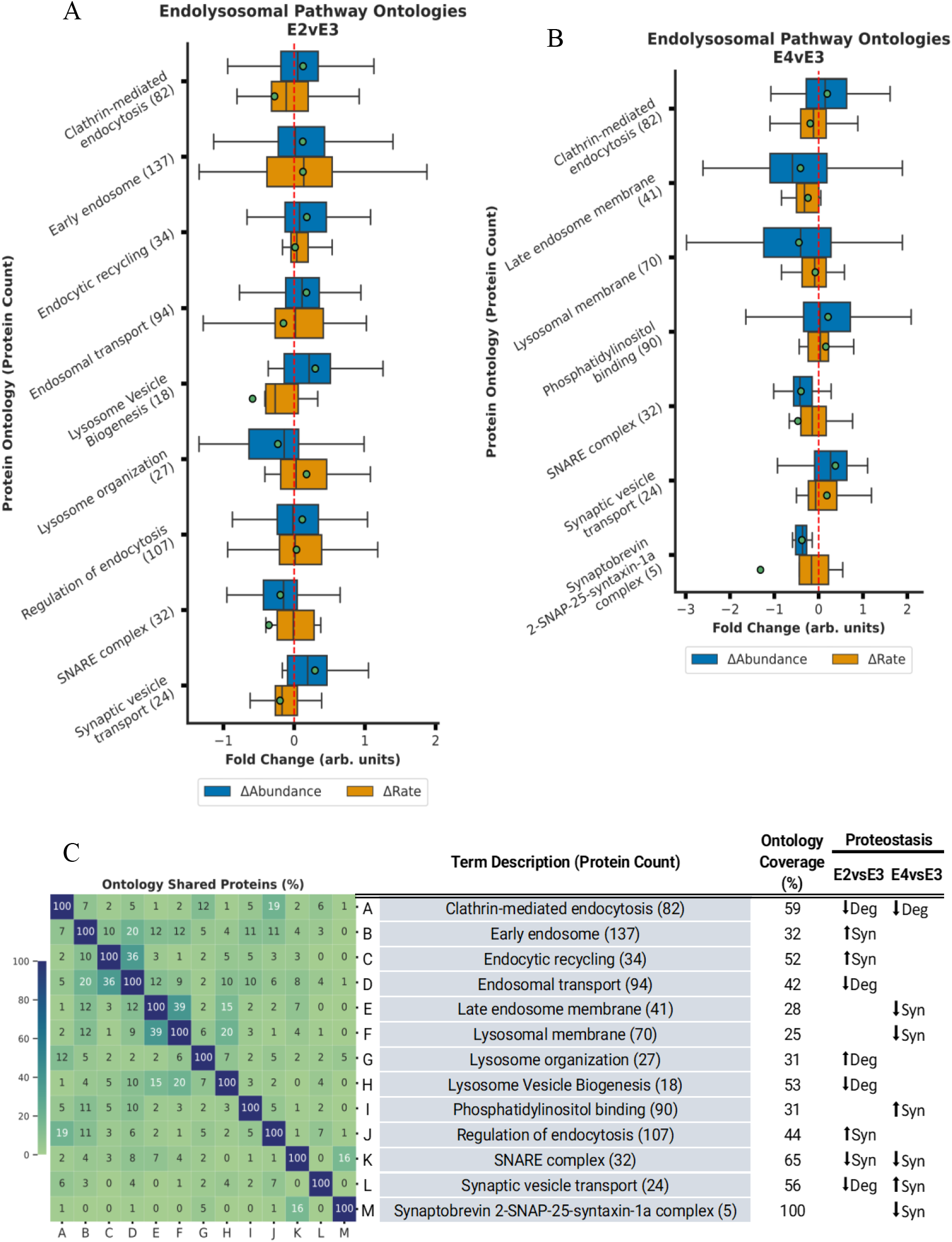
ApoE4 expression disrupts endosomal maturation and ApoE2 increases lysosomal capacity. **A-B**. Bar plot displaying Δabundance (orange) and Δturnover (blue) of proteins detected in all experimental cohorts for significant* ontologies related to endolysosomal trafficking in the E2vsE3 comparison (A) and in the E4vsE3 comparison (B). **C**. Heatmap displaying % of proteins shared across the endolysosomal ontologies with significant* Δabundance. *(*BH-PV < 0*.*05)*.

### ApoE Isoforms Modulate Synthesis and Degradation of Endocytic Vesicle Components

We observed that multiple ontologies with significant Δabundance were associated with endocytosis and vesicular processing (Figure 2). Specifically, the general *Endocytosis (158)* ontology demonstrated increased Δabundance and decreased Δturnover, suggesting reduced degradation in both ApoE2 and ApoE4 compared to ApoE3. In the context of ApoE2, *Clathrin-mediated endocytosis (82), Clathrin binding (35*), and *Clathrin coat (26)* mirrored the same ↓degradation effect observed in endocytosis, while *SNARE complex (32)* showed diminished Δabundance and Δturnover, suggesting a decline in protein synthesis compared to ApoE3. Moreover, ApoE2 expression led to significant alterations in several regulatory ontologies tied to endocytosis and vesicular processes, such as: *Endocytic recycling (34)* (↑synthesis), *Early endosome* (↑synthesis), and *Regulation of endocytosis (15)* (↑synthesis). In ApoE2, proteins related to *Lysosome Vesicle Biogenesis (18)* have lower degradation while *Regulation of Endocytosis (107)* had increased synthesis leading to higher abundance of these protein groups and presumably more efficient endolysosomal function.

In both ApoE2 (E2vsE3) and ApoE4 (E4vsE3) we noted diminished degradation of general *lysosome (146)* proteins. Within this general ontology, the *lysosomal membrane (70)* ontology had diminished Δabundance and Δturnover only in the ApoE4 group, suggesting less synthesis of the membrane components compared to ApoE3. This is consistent with large lysosomal vesicles stored in ApoE4 cells^59, 60^. In ApoE4 mice there was higher Δabundance and Δturnover (↑synthesis) of *Phosphatidylinositol binding (90)* relative to ApoE3. Conversely, there was a decline in both Δabundance and Δturnover (↓synthesis) of *SNARE interactions in vesicular transport (19), Synaptobrevin 2-SNAP-25-syntaxin-1a complex (5)*, and *SNARE complex (32)*.

### ApoE Isoforms Modulate Synthesis and Degradation of Mitochondrial Components

Our analysis identified significant Δabundance (BH-PV < 0.05) changes for multiple ontologies related to mitochondrial components (Figure 3). In the E4vsE3 comparison, these ontologies included mitochondrial membranes, protein transport, and morphology (Figure 3A and Figure S3A). Each of these ontologies displayed a negative Δabundance coupled with a positive Δturnover, signifying ↑degradation. We also detected ↓synthesis of *mitochondrial calcium ion transmembrane transport (12)* and *mitophagy (18)*. In contrast, ApoE4 *mitochondrial matrix (159)* also had ↑degradation. (Figure 3B and Figure S3B) The percentage of overlapping proteins in each mitochondrial component ontology is displayed in Figure 3C. The key finding from these ontologies is that within the ApoE2 mice there is a coherent increase in the degradation of all mitochondrial components consistent with an increase in mitochondrial degradation as an entire unit. In contrast, the ApoE4 tissues show discordant changes in matrix versus membrane proteins suggesting that mitochondrial maintenance is more piecemeal and that mitophagy may be less efficient as previously suggested in the literature ^61^. Both the ApoE2 and the ApoE4 results are synergistic with the changes in lysosome dynamics discussed above. In ApoE2 more efficient lysosomal processing will facilitate mitophagy based quality control while the inhibited lysosomal processing would inhibit mitophagy and make the ApoE4 more reliant upon individual protein replacement strategies.

**Figure 3.**
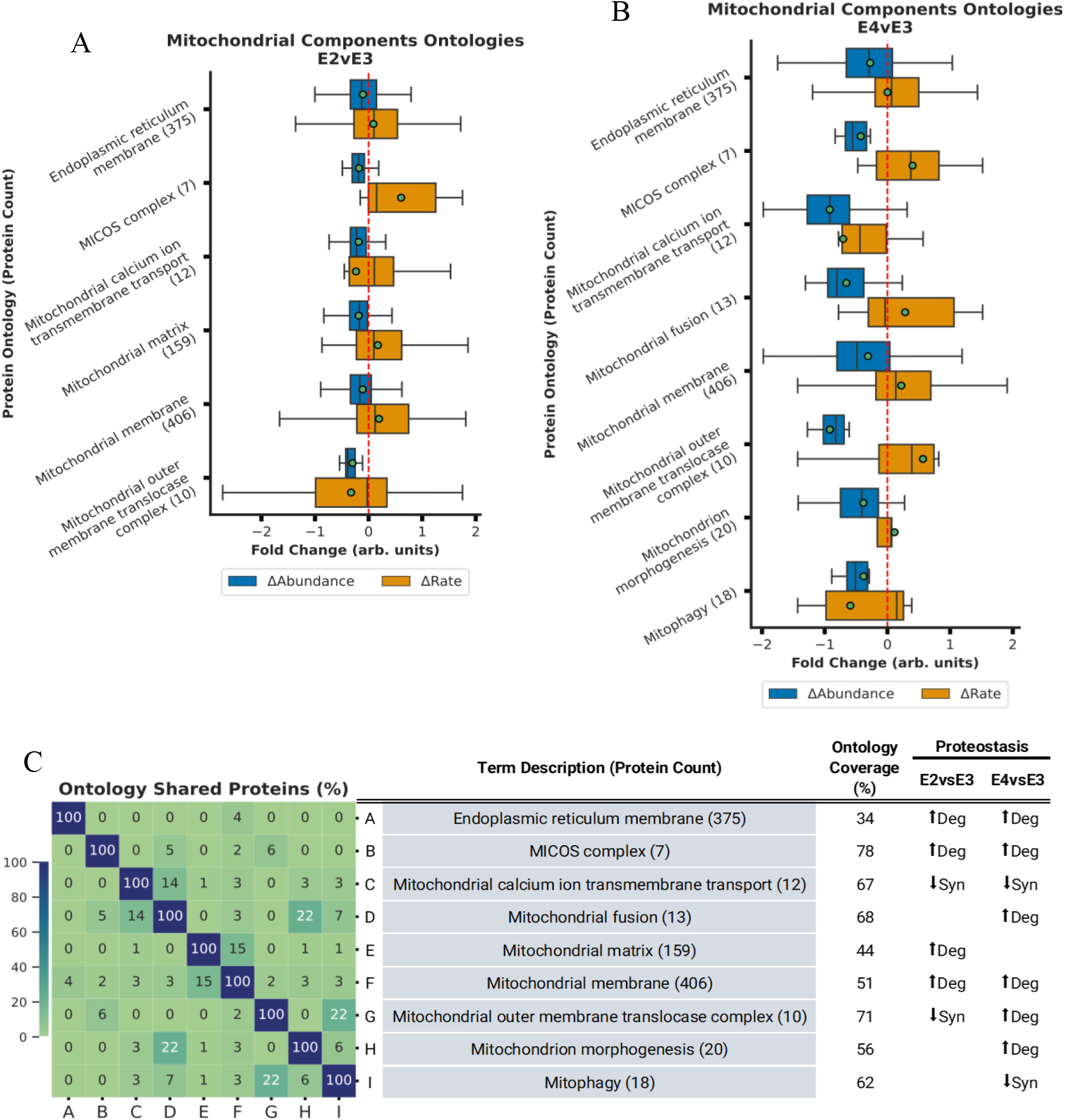
ApoE genotype differentially regulates mitochondrial proteostasis. **A-B**. Bar plot displaying Δabundance (orange) and Δturnover (blue) for ontologies of proteins detected in all experimental cohorts related to mitochondrial components with significant* Δabundance in the E2vsE3 comparison (A) and in the E4vsE3 comparison (B). **C**. Heatmap displaying % of proteins shared across the mitochondrial ontologies with significant* Δabundance. ***(*BH-PV < 0.05)***.

### ApoE4 Disrupts Metabolic Pathway Control

We observed significant changes in Δabundance (BH-PV < 0.05) across multiple ontologies related to energy production (Figure 4). ApoE2 resulted in lower expression of levels of *Pyruvate metabolism (32), Citrate cycle (TCA cycle) (26)*, and *Glycolysis/Gluconeogenesis (41)*. These reductions were primarily attributed to decreased synthesis (↓synthesis), a trend that was also evident in the *Oxidative stress and redox pathway (48)* proteins which protect the cell from reduced oxygen species. Notably, *Fatty acid beta-oxidation (25)* demonstrated reduced Δabundance coupled with increased Δturnover (↑degradation) in ApoE2 (E2vsE3), suggesting a potential decrease in fatty acid catabolism and an increase in the use of fatty acids for building complex lipids. In contrast, in the ApoE4 mice, major energy production pathways such as *Fructose and mannose metabolism (21), Pyruvate metabolism (32)*, and *Glycogen metabolism (22)* all exhibited increased Δabundance and Δturnover, pointing towards enhanced synthesis of enzymes involved in carbohydrate metabolism in the E4vsE3 comparison and an increased reliance on carbohydrates for energy similar to previous observations ^62, 63^.

**Figure 4.**
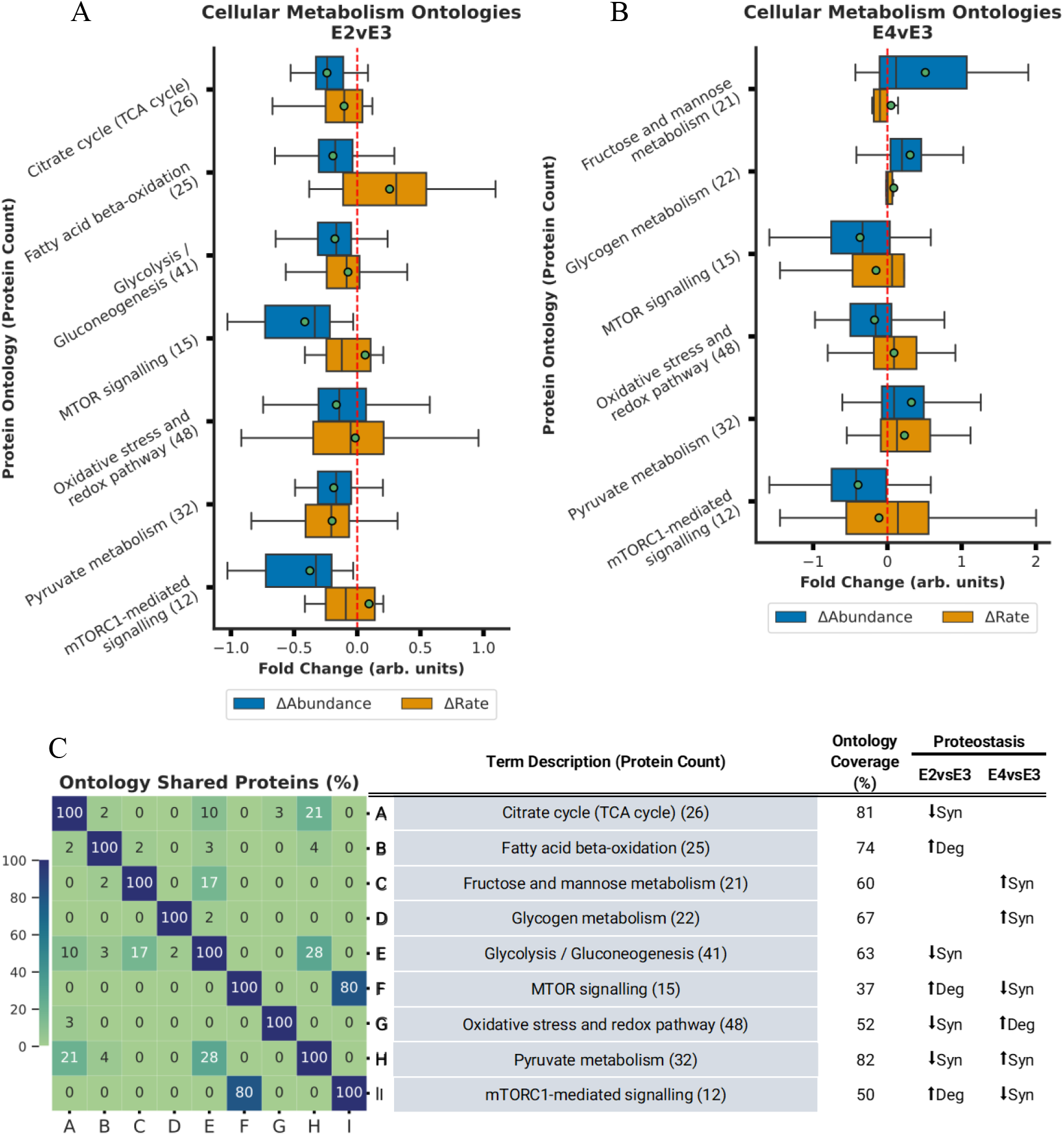
ApoE2 and ApoE4 expression drive changes in cellular fuel selection **A-B**. Bar plot displaying Δabundance (orange) and Δturnover (blue) for ontologies of proteins detected in all experimental cohorts belonging to oxidative phosphorylation with significant* Δabundance in the E2vsE3 comparison (A) and in the E4vsE3 comparison (B). **C**. Heatmap displaying % of proteins shared across the oxidative phosphorylation ontologies with significant* Δabundance. ***(*BH-PV < 0.05)***.

### ApoE Isoforms and Ubiquitin-Proteasome Pathway Activity

The proteasome related ontologies exhibited significant changes in regulation due to ApoE isoforms (Figure 5). For both the E2vsE3 and E4vsE3 comparisons, we identified pronounced increases in Δabundance and reductions in Δturnover (↓degradation) associated with the *proteasome complex (47)* Furthermore, an increased Δabundance and Δturnover (↑synthesis) of proteins involved in the *Regulation of ubiquitin-dependent protein catabolic process (62)* was statistically significant in both comparisons (BH-PV 0.05).

**Figure 5.**
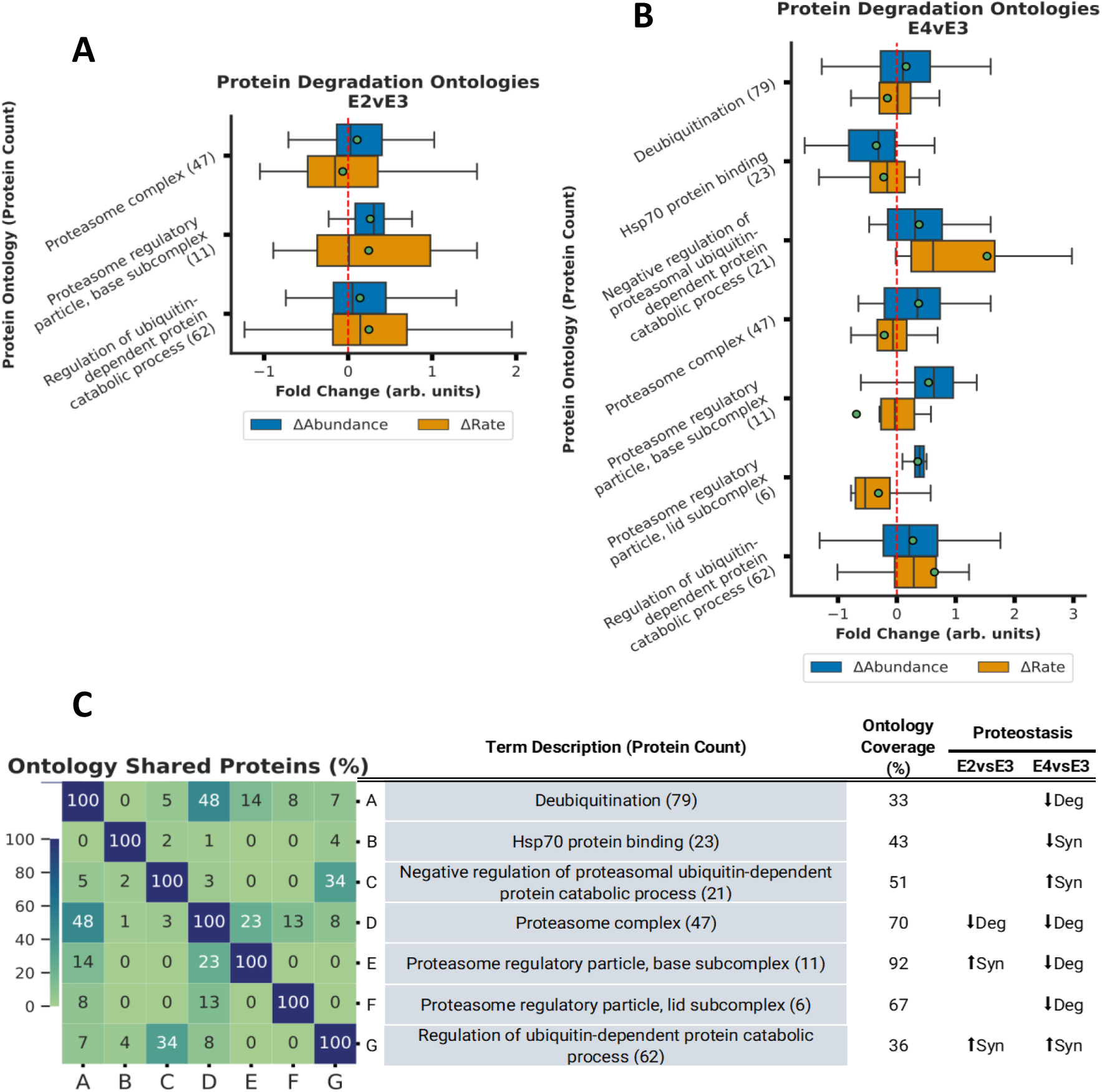
Proteasomal activity decreases with ApoE4 expression. **A-B**. Bar plot displaying Δabundance (orange) and Δturnover (blue) of proteins detected in all experimental cohorts for several ontologies related to proteasomal activity with significant* Δabundance in the E2vsE3 comparison (A) and in the E4vsE3 comparison (B). **C**. Heatmap displaying % of proteins shared across the ontologies with significant* Δabundance. ***(*BH-PV < 0.05)***.

The *proteasome regulatory particle, base subcomplex (11)* displayed ↑synthesis in the E2vsE3 comparison and ↓degradation in the E4vsE3 comparison. Meanwhile, proteins within the *Proteasome regulatory particle, lid subcomplex (6)*, demonstrated significant Δabundance due to ↓degradation in the E4vsE3 comparison with ApoE4 expression. However, these changes were not significant in the E2vsE3 comparison. Additionally, in ApoE4 we noted ↑synthesis in the *Negative regulation of proteasomal ubiquitin-dependent protein catabolic process (21)*, ↓degradation *Deubiquitination (79)*, and ↓synthesis *Hsp70 protein binding (23)*. These observations suggest a nuanced regulation of the ubiquitin-proteasome system (UPS) in association with ApoE isoforms. Hsp70 proteins are often deemed pivotal regulators of proteasome activity^64^. These changes suggest a significant reduction in the protein quality control for ApoE4 tissue (Figure 5C).

### Quantifying ApoE-dependent Shifts in Liver Proteostasis

The liver is the largest producer of ApoE in the body and is also a major receptor of ApoE and its associated cargo^9, 65, 66^. Therefore, we tested whether the liver tissue from these same experimental mice would show matching ApoE allele-specific shifts in proteome regulation.

In ApoE2 liver there was not a significant change in any of the endocytic processes relative to ApoE3 (Table S5). Multiple mitochondrial ontologies in the liver changed in significant ways and nearly 60% of their proteostasis changes are equivalent to the brain. Most changes in the mitochondria in the liver with ApoE2 expression involve increased degradation of mitochondrial components, though there is some reduced synthesis for the mitochondrial envelope and transmembrane transport. Interestingly, where ApoE2 the brain contains decreased degradation of proteasomal components, in the liver we observed increased synthesis and greater more proteasomal capacity similar to published studies^67-69^.

In ApoE4/E3 liver comparisons there was not a significant change in any of the endocytic processes (Table S5) with the exception of endosomal protein localization. In the brain, this ontology had increased synthesis, while the liver promotes decreased degradation, both of which result in an increased concentration. As for ApoE4 mitochondrial components, most changes in the liver involve increased degradation. All of the significant increased degradation ontologies observed in the brain were observed with the similar increased degradation in the liver, though not all were significant. Likewise 80% of the significant mitochondrial liver ontologies had the same proteostasis changes in the brain (Table S5). The 20% differences were due to certain NADH and ATP synthesis electron transport chain ontologies that were increased synthesis in the brain and increased degradation in the liver. Similar to the liver comparison of ApoE2 with the proteasome, in ApoE4 liver data there were no shared proteasome proteostasis changes with the brain [Figure S7 and S8]. These data suggest that most of the ApoE effects observed in the brain are not global.

## Discussion

### Exploring ApoE-genotype Effects Through the Lens of Proteostasis

Compared to the neutral ApoE3 allele, expression of ApoE4 heightens the risk for neurodegeneration, while the expression of ApoE2 is protective^6, 9, 63, 67, 70-73^. We conducted an experiment to identify how protein homeostasis changes with ApoE genotype in the tissues of human-ApoE transgenic mice (Fig 1A). Homeostasis is the dynamic control of concentration and quality in the cell (Figure S2). Traditionally, the dynamic control of protein concentration is conceptualized as the balance between synthesis and degradation^26^ while protein turnover rate is defined as the time required for a protein to be replaced.^24, 32, 58, 74^ We present this as simplified equations that relate to synthesis and degradation (Fig 1B, see ‘*Proteostasis Model and Analysis Rational*’ section for more detail). Therefore, changes in protein expression levels (Δabundance) paired with changes in protein turnover rate (Δturnover) can highlight the regulatory balance of synthesis versus degradation between conditions (Figure 1B-C).

For instance, to increase [protein] in the experimental condition (resulting in a positive Δabundance), cells can either elevate synthesis or diminish degradation. Alternatively, to decrease [protein] (leading to a negative Δabundance), cells might reduce synthesis or increase degradation. Using the change in protein turnover rates (turnover = ½(synthesis + degradation), *Proteostasis Model* section Equation 10) we can deduce whether changes in synthesis or degradation led to changes in abundance (Figure 1B). Therefore a positive Δabundance indicates increased synthesis when Δturnover is positive or reduced degradation when Δturnover is negative. Conversely, a negative Δabundance signals increased degradation when Δturnover is positive or decreased synthesis when Δturnover is negative (Figure 1C). We used this model to identify the ApoE-dependent changes in proteostasis regulation (Figure 6). Below we discuss how our results unify a diverse set of literature observations where ApoE-dependent modifications of endosome trafficking, as well as lysosomal, mitochondrial, and proteosomal function have been reported.

**Figure 6.**
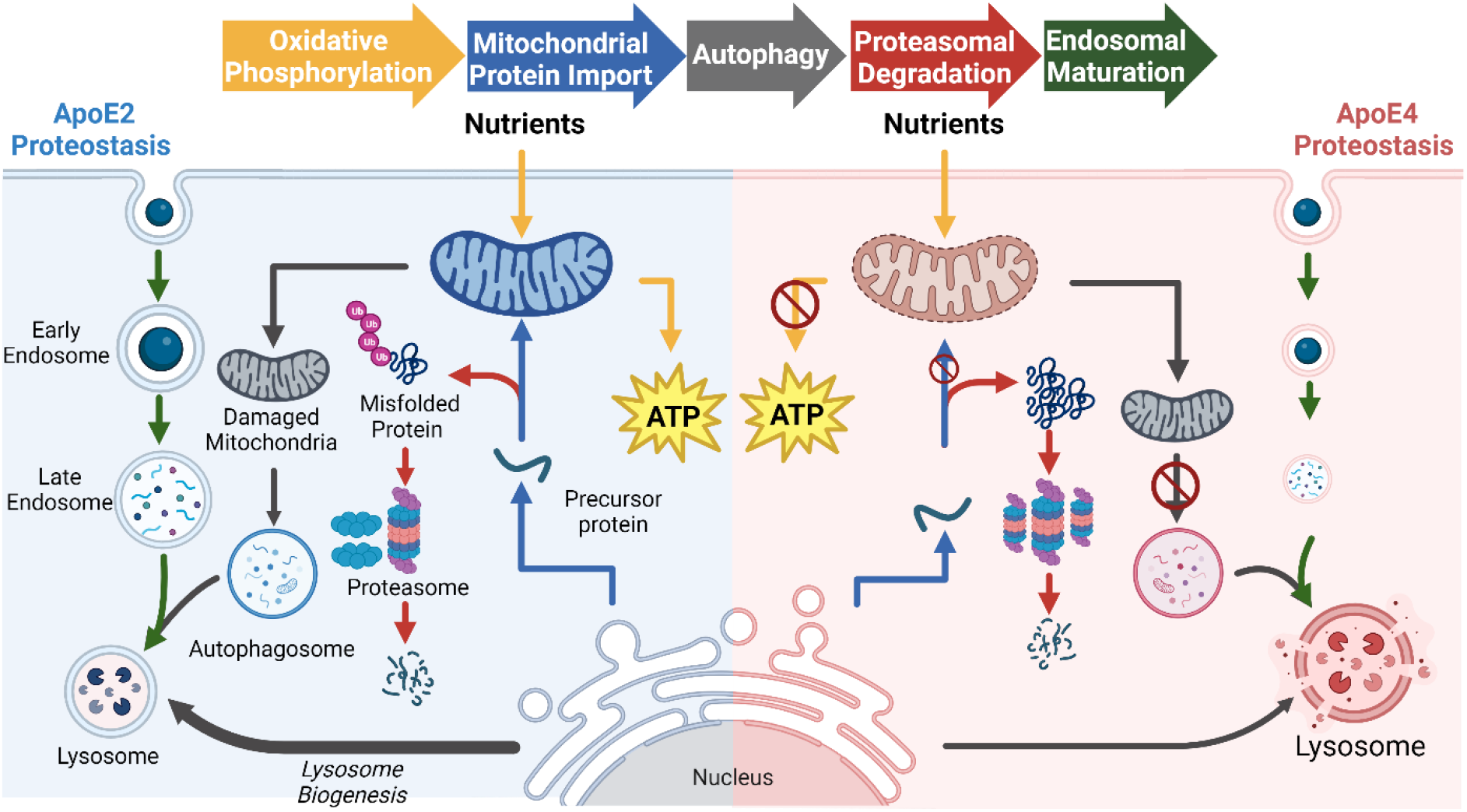
Model comparing the observed changes in proteostasis for ApoE2 and ApoE4. The arrows are color coded to represent the different pathways impacted in both ApoE2 and ApoE4 when compared to ApoE3

### ApoE isoforms modify Endocytic/Endosomal trafficking

Previous research has highlighted the dysregulation of endocytic pathways associated with ApoE4 expression^59, 60, 70, 75-77^. We detected notable ApoE4-dependent changes in several ontologies related to endocytosis (Figure 2 and Figure S3). This is in line with what is known about how ApoE isoforms modify affinity for cell surface receptors, such as LDLR and APOER2^4, 5^, initiating the endocytosis of ApoE along with its content. After this endocytic event, ApoE-laden endosomes undergo various maturation stages, wherein contents are earmarked for recycling, delivery, or degradation.

ApoE4 has a higher binding affinity to receptors^4, 5^ and is known to induce a trafficking anomaly in the early endosome^78^, then lead to accumulation and enlargement of lysoendosomal compartments^59, 60^. Following an endocytic event, the clathrin coat dismantles, allowing vesicles to transit to various destinations for cargo release. This fusion mechanism leans heavily on SNARE and SNAP-receptor proteins, also pivotal for exocytosis. In our study, we observed diminished synthesis of SNARE and SNAP ontologies in ApoE4 mice which may disrupt vesicle fusion between organelles^79^ and in response to exocytic sequences^80^. Our study revealed a reduced degradation of proteins associated with *Clathrin-mediated endocytosis (82)*, increased synthesis of *PICALM (90)*, and reduced synthesis of the *lysosomal membrane (70)* (Figure 2B, Figure S3B) in the presence of ApoE4. Priyanka et al. also noted a decline in clathrin-mediated endocytosis in astrocytes with ApoE4 expression^81^ while *in vivo* studies identified alterations in early endosome populations in 18- and 25-month-old ApoE4 mice^82^. This mechanism further incorporates phosphatidylinositol binding proteins like PICALM^83^. Before undergoing lysosomal degradation, endosomes transition to the late endosomal phase. Our findings suggest that ApoE4 expression reduces the synthesis of both late endosomal and lysosomal membranes. Although the general *lysosome (146)* ontology exhibits increased degradation with ApoE4 expression, if we look specifically at the membrane components of this ontology then the ApoE4 specifically has less total protein due to lower synthesis. These results are consistent with previous observations of large-volume lysosomes which would have a low membrane surface/volume ratios accumulating in the in ApoE4 cells ^59, 60^. Collectively, these results underscore multiple points of failure due to ApoE4-associated inhibition of endosomal maturation and stalled lysosomal functions as previously observed ^78, 82, 84^.

The E2vsE3 tissue had similarities in vesicle-centric ontologies. Notably, there was a decline in the degradation of endocytosis and clathrin protein-related ontologies, and SNARE complexes saw reduced synthesis. This implies that ApoE2 also modifies vesicle endocytosis. However, the changes suggested a more streamlined regulation of endolysosomal events with ApoE2 (E2vsE3). This again agrees with literature reports of modified receptor binding with ApoE2 having lower affinity while ApoE4 has a higher affinity^4, 5^. This coupled with lower degradation of proteins within the lysosome vesicle biogenesis ontology for the E2vsE3 comparison, a process intrinsically tied to endosomal trafficking and central to lysosomal adaptation^85^, suggests a tighter control of endocytic events and better lysosomal quality with ApoE2 expression. These observations agree with previous research on astrocytes indicating ApoE2 expression increases lysosomal activity relative to ApoE3 and ApoE4 expression^86^.

### ApoE-dependent changes in Mitochondrial Proteostasis

We observed ApoE-dependent changes in mitochondrial proteostasis that were consistent with modified autophagy and lysosomal function. In ApoE4 (E4vsE3) mice, we measured elevated degradation in *mitochondrial membrane (406), mitochondrial inner membrane (303)* (Figure 3 and Figure S4), *Cristae formation (11), Mitochondrial fusion (13)*, and *mitochondrial transport (72)* with no accompanying change in general *mitochondrial matrix (159)* and decreased synthesis of *mitophagy (18)*. Mitochondrial membrane complexes play critical functions in cellular homeostasis—such as energy production, calcium level modulation, apoptosis, and the regulation of reactive oxygen species (ROS) ^87^. Prior research has documented alterations in the mitochondrial membrane’s integrity in the context of neurodegenerative diseases ^88, 89^.

Most mitochondrial proteins are encoded on the nuclear DNA and are transported into the mitochondria through translocases (TIM and TOM)^90, 91^. These translocases interact with the many inner mitochondrial membrane (IMM) folds that make up the cristae via the mitochondrial cristae organizing system (MICOS).^92^ Our observations indicate a change in mitochondrial protein import, especially evident in the higher degradation of the *TOM complex (10), cristae formation (11)*, and the *MICOS complex* (7) ontologies. (See Figure 5B and Figure S5B). The MICOS also plays a vital role in cristae organization and the function of respiratory complexes.^93^ Disruptions in MICOS have been documented to modify cristae structure^94^, and recent studies associate altered MICOS protein expressions with ApoE4 manifestation^89^. The MICOS literature also report evidence of mitochondrial fusion and fission imbalance in Neuro-2a cells expressing ApoE4.^89^ Previous analysis of AD brains indicated diminished protein levels connected to mitochondrial fusion/fission^95^, which our data supports as a degradation driven loss of fusion proteins (See Figure 3B and Figure S3B).

Mitochondria and the endoplasmic reticulum (ER) collectively form the mitochondria-associated membrane (MAM), which has implications in AD pathology^96-98^. These MAMs regulate oxidative phosphorylation, calcium levels, protein degradation, and mitochondrial membrane organization. Our dataset elucidates an ApoE4-induced *MAM (57)*, marked by increased degradation contrasted against ApoE3 (Figure 3 and Figure S4). Our results support ApoE4-related MAM instability by diminished synthesis of chaperone complexes, mitophagy, and calcium transport.

In contrast, ApoE2 mice display increased degradation of mitochondrial membrane ontologies with a matching increase in the degradation of the matrix proteins. Although both ApoE2 and ApoE4 mice revealed changes in the mitochondrial membrane and transport, our ApoE2 findings suggests that there is a cohesive organelle-wide response involving both membrane and matrix proteins. Drawing from our preliminary insights into endolysosomal systems discussed above, we postulate this might be evidence of superior mitophagy in ApoE2. Additionally, we theorize, as described in the literature,^61-63, 89, 99^ that the alterations in mitochondrial proteostasis induced by ApoE4 culminate in mitochondrial dysfunction.

### ApoE disrupts ATP production

There is an increasing body of research on ApoE genotype-specific effects on ATP production^100-102^. Moreover, compromised bioenergetic pathways are identified as a distinct characteristic of neurodegeneration^103-106^. Several studies highlight an ApoE-related shift towards glycolysis and diminished oxygen consumption in brain tissues^103, 107^. Our data align with these observations, revealing a heightened synthesis of ontologies suggesting that ApoE4 expression leads to a more pronounced reliance on glycolytic pathways compared to ApoE3 (see Figure 4B, Figure S4B). These ontologies include *Fructose and mannose metabolism (21), Pyruvate metabolism (32)*, and *Glycogen metabolism (22)*. We posit that this increased reliance on carbohydrate metabolism is a consequence of the lack of cohesive mitochondrial maintenance. Another study focusing on glycolytic and OXPHOS activities found that ApoE4 expression leads to compromised mitophagy and elevated ROS levels in brain cells, a trend our data supports^108^. The impaired mitophagy was attributed to cholesterol buildup in lysosomes. While we haven’t analyzed cholesterol or ROS levels, our data does indicate reduced synthesis in *mitophagy (18), lysosomal membrane (70)* and *Detoxification of Reactive Oxygen Species (20)* proteins —potential indicators of disrupted mitophagy and ROS balance (See Figure 3B, Figure S3B)

In cell culture ApoE2 expression has been associated with enhanced ATP production and heightened glycolytic activity^103, 107^. In contrast, various studies have shown that ApoE4 expression was associated with diminished ATP production^100, 109-111^. In ApoE2 tissue we observed decreased abundance in ontologies such as the *TCA cycle (25), Pyruvate metabolism (32)*, and *Glycolysis / Gluconeogenesis (41)* due to diminished synthesis and augmented degradation. Our study is averaging together all cell types in the brain and therefore may diverge from cell type-specific experiments ^100^. Collectively, our data accentuates the isoform-specific alterations in diverse metabolic pathways, and suggests that isolating single cell types from the brain may be a critical method to test for metabolic changes in response to ApoE isoforms.

### Linkages between the Proteasome and Mitochondrial Homeostasis

The proteasome is a key component of proteostasis maintenance and is essential in clearing out misfolded proteins and saving cells from misfolding stress response.^112^ Reduced proteasome activity has consistently been implicated as a major player in the pathophysiology of neurodegeneration^23, 61, 67, 69, 113^. ApoE has been shown to directly regulate the cleavage of amyloid precursor protein (APP) to form amyloidogenic Aβ peptides with ApoE4 allowing increased Aβ peptide cleavage and plaque deposition.^114^ This buildup of Aβ is relevant to proteasomal function and has been shown to directly inhibit proteasome function leading to increased accumulation of amyloid plaques.^115, 116^

The proteasome also plays a major role in the mitochondrial quality control, especially in response to misfolded proteins that disrupt mitochondrial activity^117^. Interestingly, a growing body of literature suggests that proteasome function can also be disrupted by mitochondrial dysfunction. For example, oxidation of the 26S subunit of the proteasome due to increased mitochondrial oxidative stress has been shown to increase the accumulation of ubiquitinated substrates and decrease proteasomal activity^118^. Notably, we discerned a significant reduction in the synthesis of HSP70 proteins in E4 which latch onto misfolded or compromised proteins before proteasomal degradation^68^. This interconnection of the proteasome and mitochondria as well as their consistent implication in neurodegenerative disease has led some researchers to suggest that dysfunction in either the proteasome or mitochondria are “two sides of the same coin” leading to a futile cycle of mitochondrial and proteasomal insult.^119^

Our investigation revealed reduced degradation of both the proteasome and its regulatory complex in ApoE4 mice (refer to Figure 5 and Figure S6). Additionally, while ApoE4 expression increased the synthesis of ontologies linked to ubiquitin-proteasome regulation, it also elevated the synthesis specifically for its negative regulation. Such trends align with existing literature delineating the impact of ApoE isoforms on proteasomal dysregulation in Alzheimer’s disease^67, 120^. Alongside our observation of compromised mitochondrial activity in ApoE4 mice, these findings imply a heightened susceptibility to both mitochondrial and proteasomal damage. The concurrent malfunction of mitochondria and proteasomes has been historically correlated with neuronal apoptotic pathways and neurodegeneration, thereby underlining a mechanism through which ApoE4 exacerbates the risk of neurodegenerative ailments such as Alzheimer’s^121^.

ApoE2 also exhibited a notable decline in the degradation of proteins associated with ubiquitin and proteasomal processes, paralleling the ApoE4 response (see Figure 5 and Figure S6). While both ApoE2 and ApoE4 amplify the regulation of ubiquitin-dependent catabolism, the base complex of the proteasome (responsible for facilitating the unfolding and admission of ubiquitinated polypeptides into the proteasome’s degradation chamber^122^) reduced degradation of deubiquitinating proteins was more prevalent in ApoE4. This implies larger proteasome pool in ApoE4, albeit with more regulation. It’s worth speculating that the ApoE2-mediated inhibition of APP cleavage to form Aβ^114^ might also be instrumental in curbing proteasomal dysfunction.

These patterns suggest that both phenomena might be interconnected, with proteasomal dysfunction potentially instigating ApoE4-associated reductions in ATP production. Beyond the ubiquitin-proteasome system, autophagy is instrumental in clearing defective mitochondria via lysosomal degradation. As previously mentioned, our findings substantiate the ApoE4-associated dysregulation of MAM structures, which potentially results in disrupted mitochondrial morphology and impaired energy production. Our working hypothesis postulates that ApoE4 expression precipitates a decline in proteasomal activity, culminating in the accrual of dysfunctional mitochondria and a diminished capacity to eliminate these via autophagy. Drawing from our data on lysosomal components in ApoE4 and existing literature^78, 123^, we surmise that suboptimal endocytic regulation might directly impact autophagy and the proteasomal oversight of mitochondrial proteostasis. Conversely, ApoE2 expression is purportedly linked to enhanced proteasomal capability and autophagy through lysosomal degradation, resulting in fortified mitochondrial proteostasis. Our objective is to delve deeper into these discoveries and authenticate our hypothesis across various ApoE models.

### Liver Proteostasis Changes Compared to Brain

ApoE is an important lipid transporter that is expressed and integral to many parts of the body beyondthe brain. Previous literature has shown that despite ApoE2’s protective effect against Alzheimer’s Disease, it increases risk for cardiovascular health^66^. This leads us to question whether the mechanism by which ApoE2 protects against Alzheimer’s Disease may in turn be detrimental and disease causing to other tissue. To determine whether ApoE does indeed elicit a global response across tissue, we tested and analyzed the liver tissue from our experimental mice in the same manner as the brain.

The changes in ApoE2/E3 liver proteostasis and ApoE2/E3 brain proteostasis were not equivalent. Although many trends were similar between brain and liver, few changes were statistically significant in the liver. Mitochondrial protein localization, transportation, and organization had shared proteostasis changes between tissues, but general trends from other ontological changes suggest mitochondrion turnover to be reduced in the liver compared to the brain (See Figure S2).

ApoE4/E3 liver and ApoE4/E3 brain proteostasis changes were also not the same. This is principally due to significant proteostasis changes among the endosome, metabolism, and proteasome pathways in the brain, and that most of which were not significant in the liver (see table S5). However, comparisons between the ApoE4 liver and brain contained several of the same proteostasis changes for mitochondria, suggesting there may be some degree of shared effects due to ApoE4.

Thus, while some cellular pathways may be affected similarly, a global effect specific to ApoE allele is not supported by our data. We propose the lack of a global ApoE effect is because most tissues have a large number of apolipoproteins participating in lipid transport ^9, 66^. The brain has limited apolipoproteins compared to other tissues, in that it is limited to only the apolipoproteins it creates (primarily ApoE and ApoJ). This is due to the blood-brain-barrier, which prevents apolipoprotein transfer between the brain and the rest of the body. Since lipid trafficking in the liver has access to multiple apoliproproteins we posit that this may dilute the effect of ApoE isoforms on the pathways within liver tissue.

## CONCLUSION

In this study, we demonstrated how combining protein abundance and turnover rate unveils novel insights into the cellular mechanisms governing synthesis and degradation. Utilizing multifaceted proteomics data, we tracked ontology-level variations among distinct ApoE genotypes in healthy adult mice. Our findings present *in vivo* evidence that harmonizes with existing literature, identifying how ApoE4 interrupts endosomal trafficking leading to autophagy and proteasome activity defects and lower mitochondrial quality in the brain. Concurrently, our data suggests that ApoE2 enhances brain mitochondrial health by amplifying turnover in the brain (Figure 6).

## Supporting information

All_Supplemental_Files

## ASSOCIATED CONTENT

### SUPPORTING INFORMATION

Figure S1. Detailed workflow chart describing both the mouse model and stages of analysis.

Figure S2: Kinetic model of protein homeostasis identifying the common sources and sinks of protein in a cell

Figure S3. Abundance and turnover FCs for ontologies related to endolysosomal processes in A) E2vsE3 and B) E4vsE3.

Figure S4. Abundance and turnover FCs for ontologies related to mitochondrial components in A) E2vsE3 and B) E4vsE3.

Figure S5. Abundance and turnover FCs for ontologies related to cellular metabolism in A) E2vsE3 and B) E4vsE3.

Figure S6. Abundance and turnover FCs for ontologies rela ted to protein degradation in A) E2vsE3 and B) E4vsE3.

Figure S7. Model for comparison of the ApoE2 brain and liver homeostasis shifts relative to ApoE3. The ApoE2 brain model (left, blue), is the same as in Figure 6. The green model (right) shows the modifications in ApoE2 liver tissue.

Figure S8. Model for comparison of the ApoE4 brain and liver homeostasis shifts relative to ApoE3. The ApoE4 brain model (right, red), is the same as in Figure 6. The green model (left) shows the modifications in ApoE4 liver tissue.

The data generated in this investigation can be accessed via the ProteomeXchange Consortium via the PRIDE partner repository^124^ (http://www.proteomexchange.org/) with the accession number PXD044460. The repository includes the raw LC-MS files used for both quantitative and kinetic files used in data analysis. In addition, Peaks Studio (Bioinformics Solutions Inc.) outputs containing peptide and protein level identification data for both quantitative and kinetic measurements are included in the repository.

*Project Name: Quantitative and Kinetic Proteomics Reveal ApoE Isoform-dependent Proteostasis Adaptations in Mouse Brain*

*Project accession: PXD044460*

The output files from Deuterater software including turnover rate values are found within the repository while the code is found here (https://github.com/JC-Price/DeuteRater/releases). Lastly, the code used in the ontology analysis can be found by following this link to the GitHub repository (https://github.com/natepine/ApoE_Proteomics.git).

The following files are available free of charge via the Internet

.xls file containing Tables S1-S5 and data for all quantifiable proteins and measurable ontologies.

.pdf file containing supplementary figures S1-S7, including a diagram that explains the mouse model in addition to box plots with additional ontologies.

## AUTHOR INFORMATION

### Author Contributions

This manuscript was written through contributions of all authors. All authors have given approval to the final version of the manuscript. A CRediT statement is provided below

**Nathan R. Zuniga**: Conceptualization, Data Curation, Formal Analysis, Investigation, Methodology, Project Administration, Resources, Software, Supervision, Validation, Visualization, Writing - Original Draft, Writing - Review & Editing. **Noah E. Earls**: Data Curation, Formal Analysis, Investigation, Validation, Visualization, Writing-Original Draft, Writing - Review Editing. **Ariel E. A. Denos:** Data Curation, Formal Analysis, Supervision, Validation, Visualization, Writing – Original Draft, Writing – Review & Editing. **Jared M. Elison:** Formal Analysis, Investigation, Validation, Visualization, Writing – Original Draft, Writing – Review & Editing. **Benjamin S. Jones:** Formal Analysis, Investigation, Validation, Writing - Review & Editing. **Ethan G. Smith:** Formal Analysis, Investigation, Validation. **Noah G. Moran:** Investigation, Validation, Writing - Review & Editing. **Katie L. Brown** Investigation, Validation, Writing - Review & Editing. **Gerome M. Romero:** Data Curation, Formal Analysis, Investigation. **Chad D. Hyer:** Formal Analysis, Writing - Original Draft. **Kimberly B. Wagstaff:** Investigation, Validation. **Haifa M. Almughamsi:** Methodology, Investigation. **Mark K. Transtrum:** Methodology, Writing – Review & Editing. **John C. Price:** Conceptualization, Funding Acquisition, Methodology, Project Administration, Resources, Supervision, Writing – Original Draft, Writing - Review & Editing.

### Funding Sources

This work was made possible by a grant from the Fritz B. Burns Foundation to John C. Price; the National Institutes of Health [R01AG066874] to John C. Price; Brigham Young University Undergraduate Research Awards to Chad Hyer, Noah Earls, Noah Moran, Ethan Smith, Katie Brown, Kimberly Wagstaff, Jared Elison, and Benjamin Jones; A grant from Deanship of Scientific Research, Taif University to Haifa Almughamsi.

### Notes

The authors declare no competing financial interest.

## ACKNOWLEDGMENT

The mice used in this research were provided by Johnathan J. Wisco, Ph. D. We gratefully acknowledge the services provided by BYU’s live animal facility and the Fritz B. Burns Biological Mass Spectrometry Facility.

## ABBREVIATIONS

AD: (Alzheimer’s Disease)
Aβ: (Amyloidβ)
ApoE: (Apolipoprotein E)
APP: (Amyloid Precursor Protein)
BCA: (Bicinchoninic Acid)
BH-PV: (Benjamini-Hochberg P-value)
CID: (Collision induced dissociation)
DDA: (Data dependent acquisition)
E2/E3/E4: (ApoE2/ApoE3/ApoE4)
ETC: (Electron Transport Chain)
FC: (Fold Change)
FDR: (False discovery rate)
GFAP: (Glial fibrillary acidic protein)
IACUC: (Institutional Animal Care and Use Committee)
IP: (Intraperitoneal)
LDL: (Low density lipoprotein)
LFQ: (Label free quantification)
MS: (Mass-Spectrometry)
OXPHOS: (Oxidative Phosphorylation)
PDH: (Pyruvate dehydrogenase)
TCA: (Tricarboxylic Acid)
TIM: (mitochondrial inner membrane translocase)
TOM: (mitochondrial outer membrane translocase).

---

**Figure S2:**
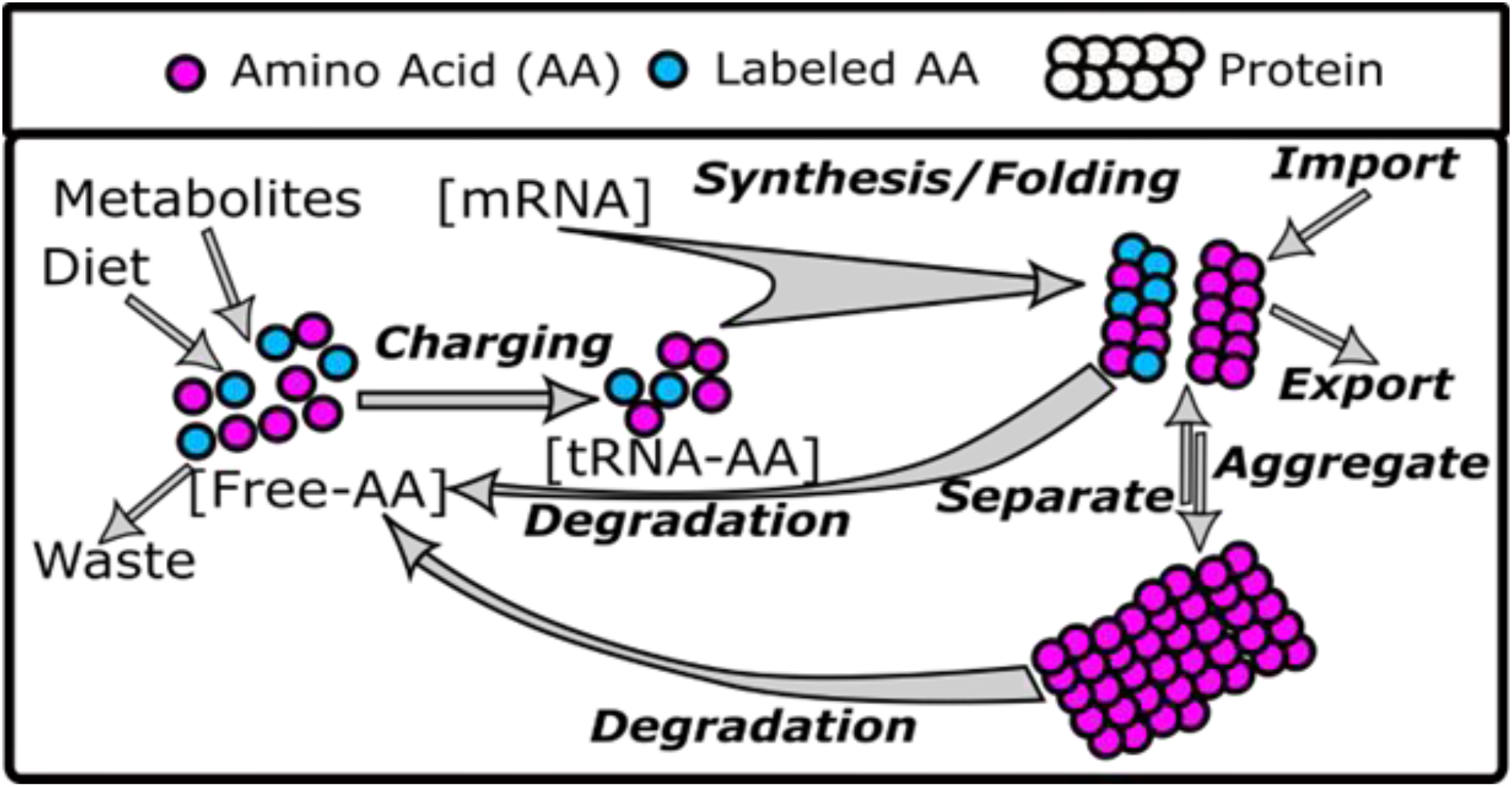
Protein Homeostasis Model with commonly observed sources and sinks of protein

## Notes

http://www.proteomexchange.org/

### Competing Interest Statement

The authors have declared no competing interest.

### Summary of Updates

Figure 1 revised, supplemental files uploaded

