## Supplementary material for "Quantitative and Kinetic Proteomics Reveal ApoE Isoform-dependent Proteostasis Adaptations in Mouse Brain": All_Supplemental_Files: Zuniga trApoE Supplementary Figures.docx

*Supplementary Figures: The box plots in the supplementary figures include additional ontologies that further support the findings of ApoE-related effects in the brains of transgenic mice. The boxplots include outliers that were not present in the main text figures because of space limitations.*

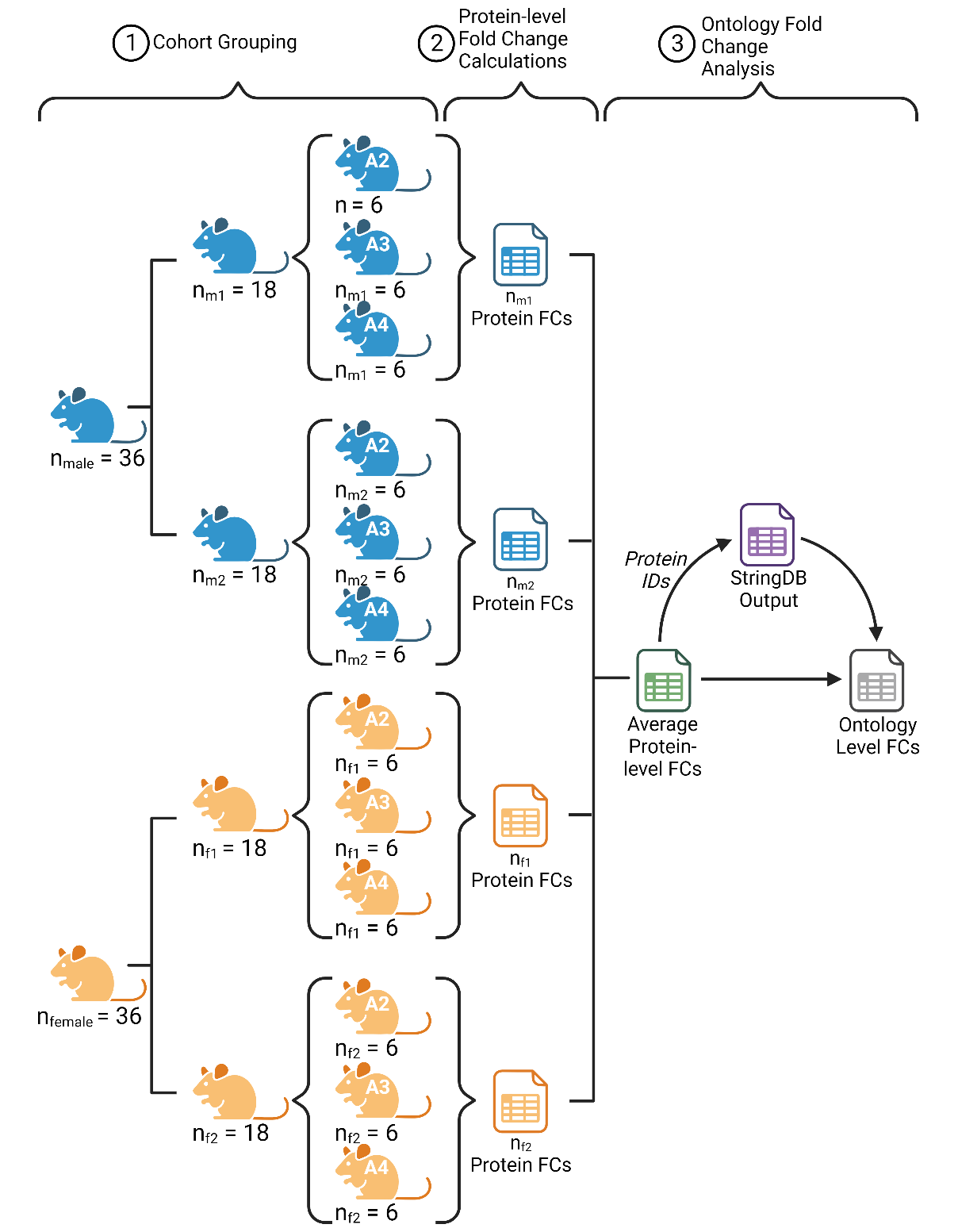

***Figure S1: ApoE Cohort Design***

*72 mice were used to explore ApoE genotype. To facilitate sample preparation and accommodate for instrument availability, mice were split into 2 male groups and 2 female groups containing a total of 18 mice in each group. Each group consisted of 6 homozygous mice of each ApoE genotype. Data from each group was analyzed independently to calculate abundance and turnover fold changes. Proteins with a quantified abundance fold change were analyzed with StringDB multiprotein tool to identify ontologies represented in those proteins. Finally, ontology level fold changes were calculated using protein fold changes from abundance and turnover data.*

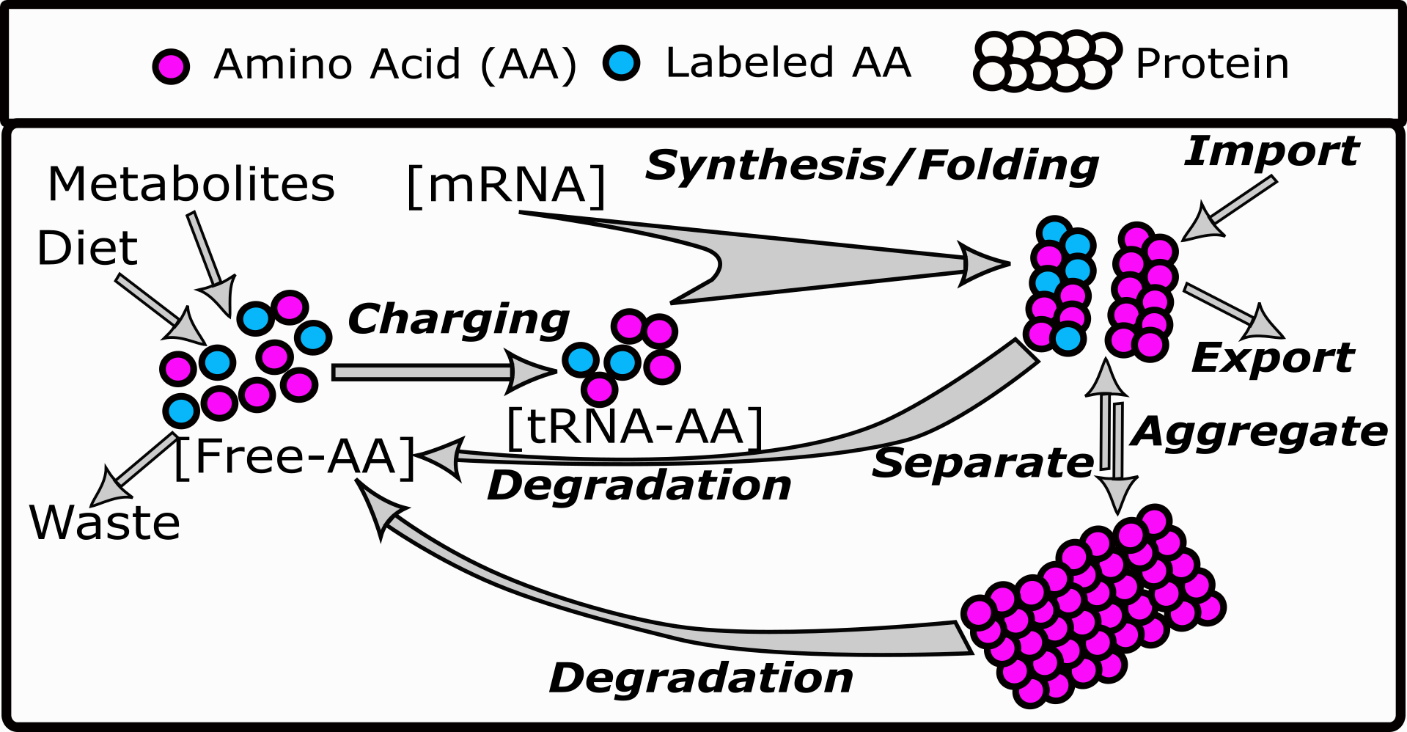

**Figure S2**: Protein Homeostasis Model with commonly observed sources and sinks of protein

 
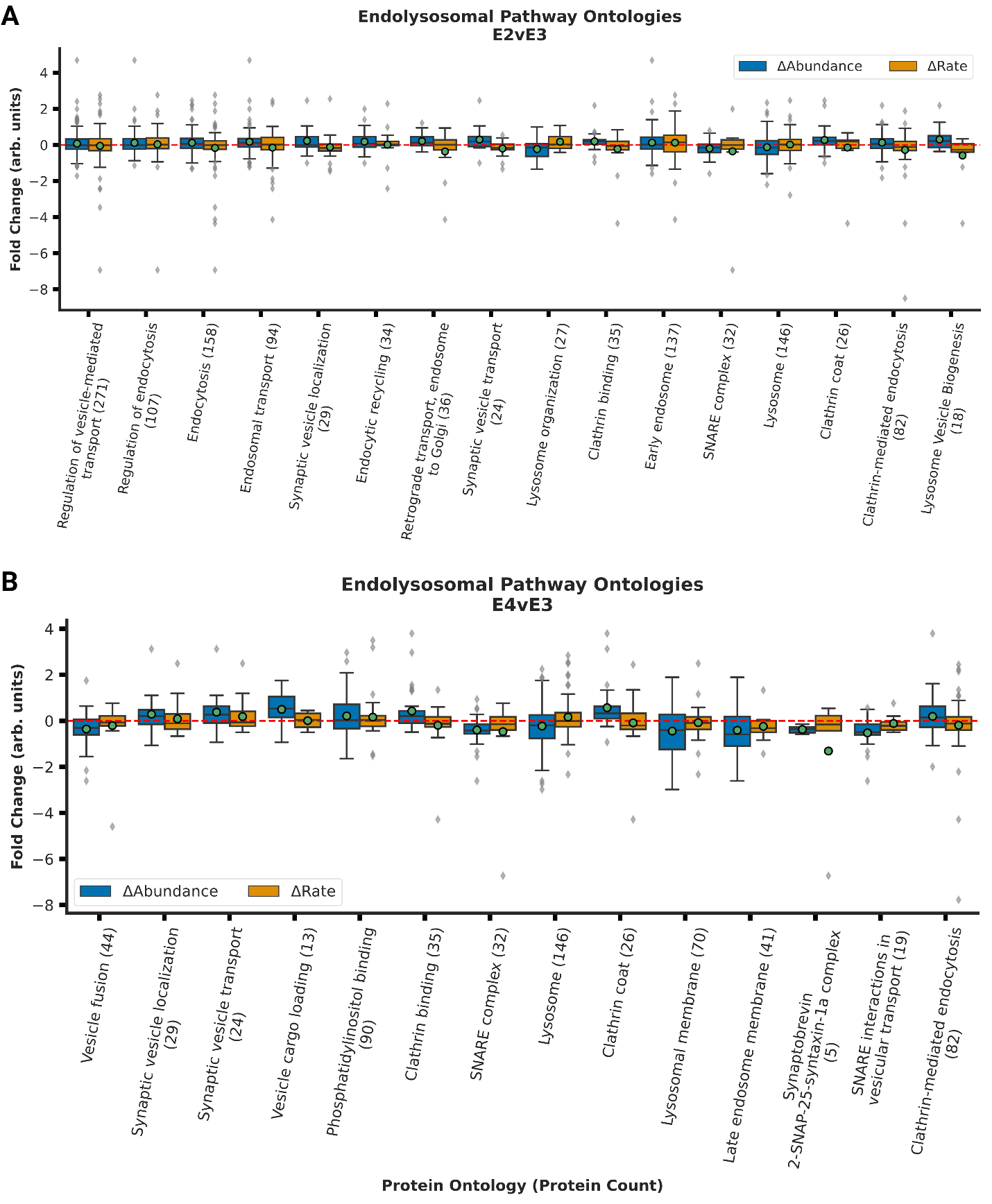

Supplementary Figure 2: Endolysosomal Ontologies

*Abundance and turnover FCs for ontologies related to endolysosomal processes in A) E2vsE3 and B) E4vsE3.*

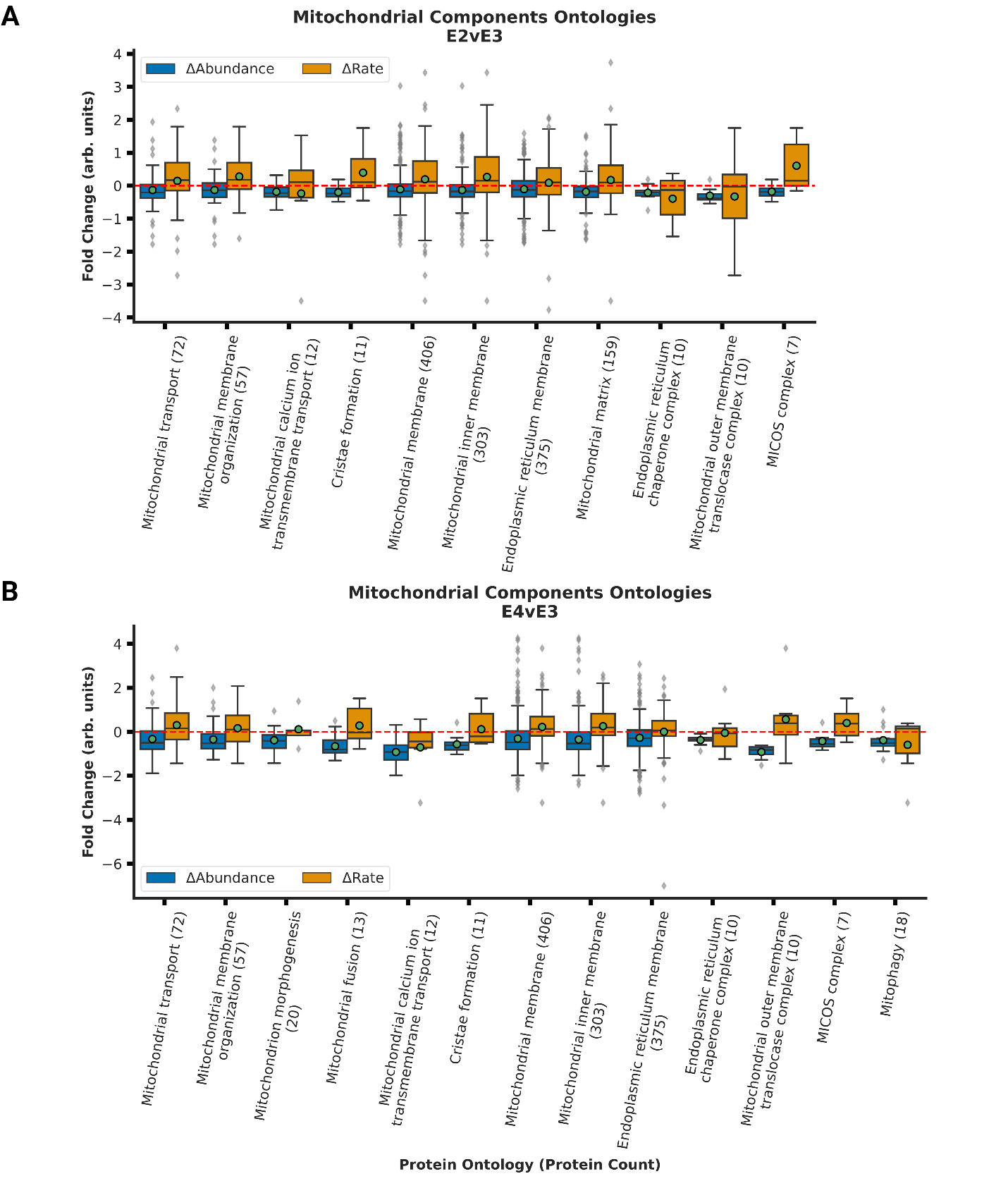

Supplementary Figure 3: Mitochondrial ontologies:

*Abundance and turnover FCs for ontologies related to mitochondrial components in A) E2vsE3 and B) E4vsE3.*

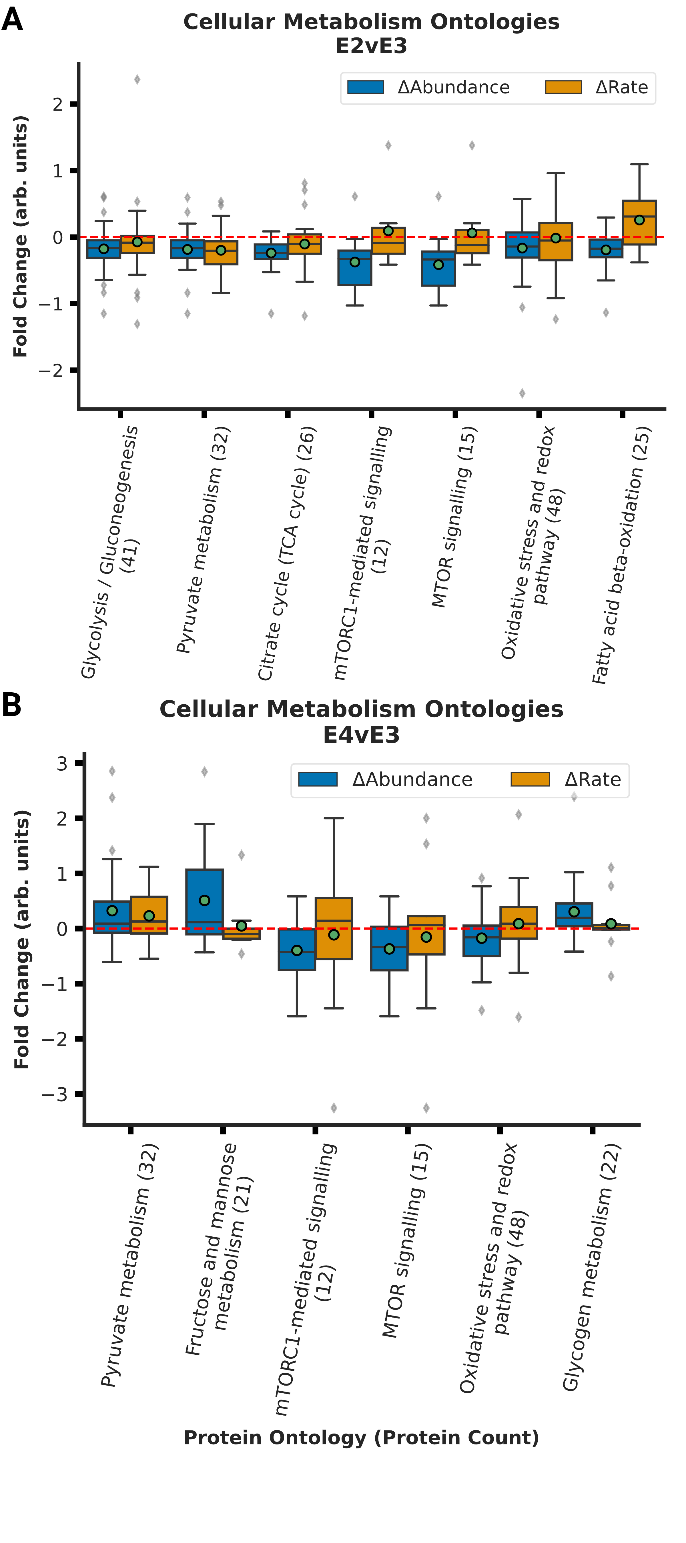

Supplementary Figure 4: Cellular Metabolism Ontologies

*Abundance and turnover FCs for ontologies related to cellular metabolism in A) E2vsE3 and B) E4vsE3.*

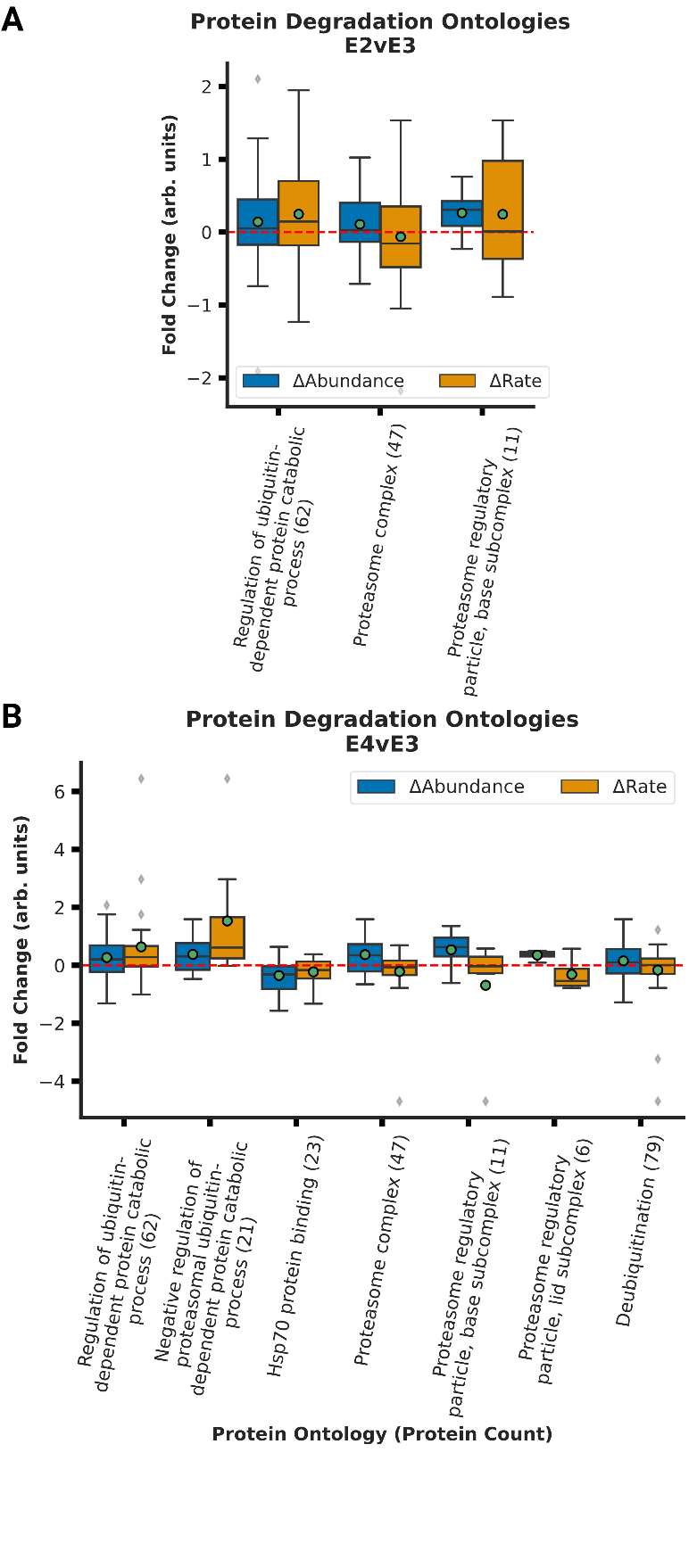

Supplementary Figure 5: Protein Degradation Ontologies

*Abundance and turnover FCs for ontologies related to protein degradation in A) E2vsE3 and B) E4vsE3.*

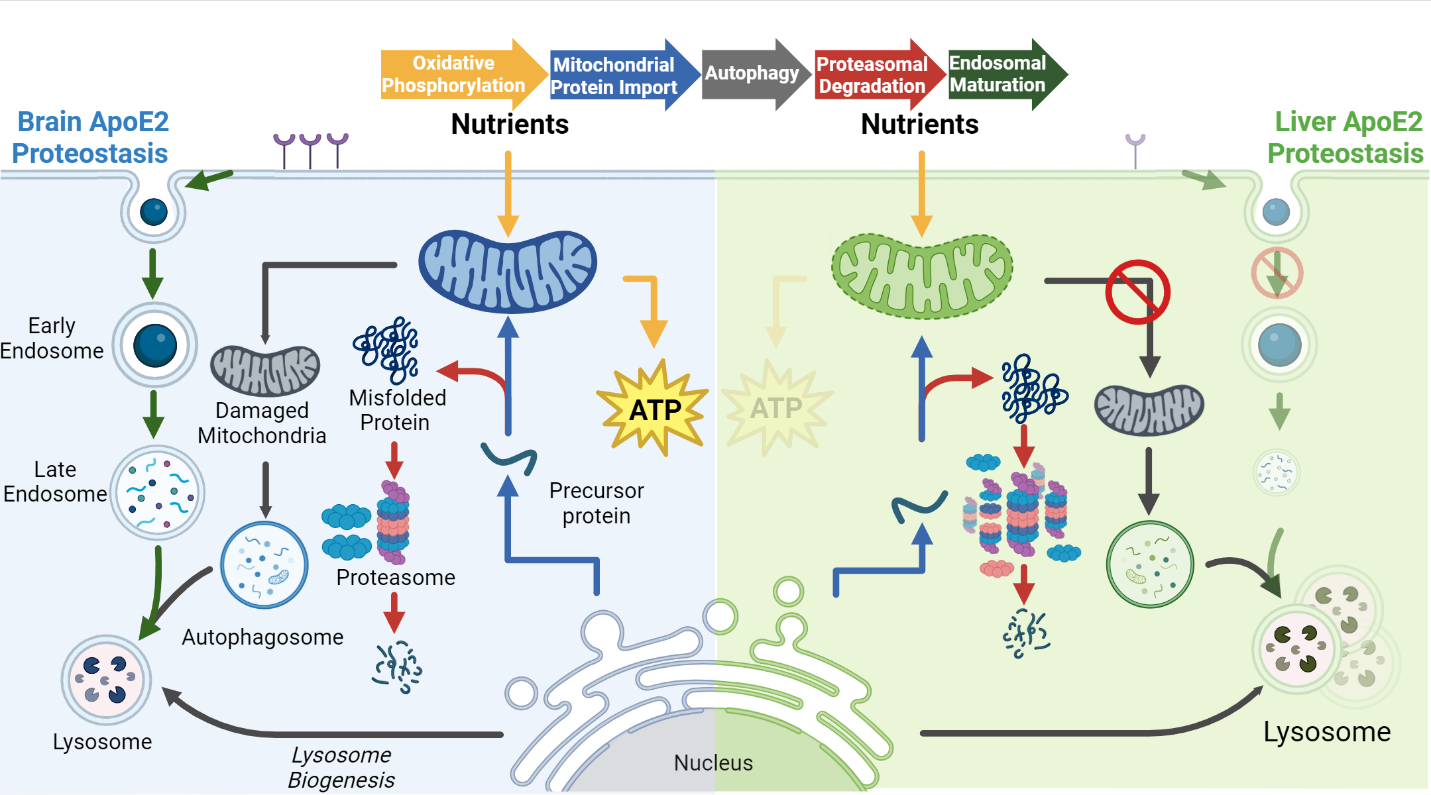

Supplementary Figure S6: ApoE2 change in brain (blue) versus liver (green) Model comparing the observed changes in proteostasis for ApoE2. The arrows are color coded to represent the different pathways impacted in both ApoE2 when compared to ApoE3

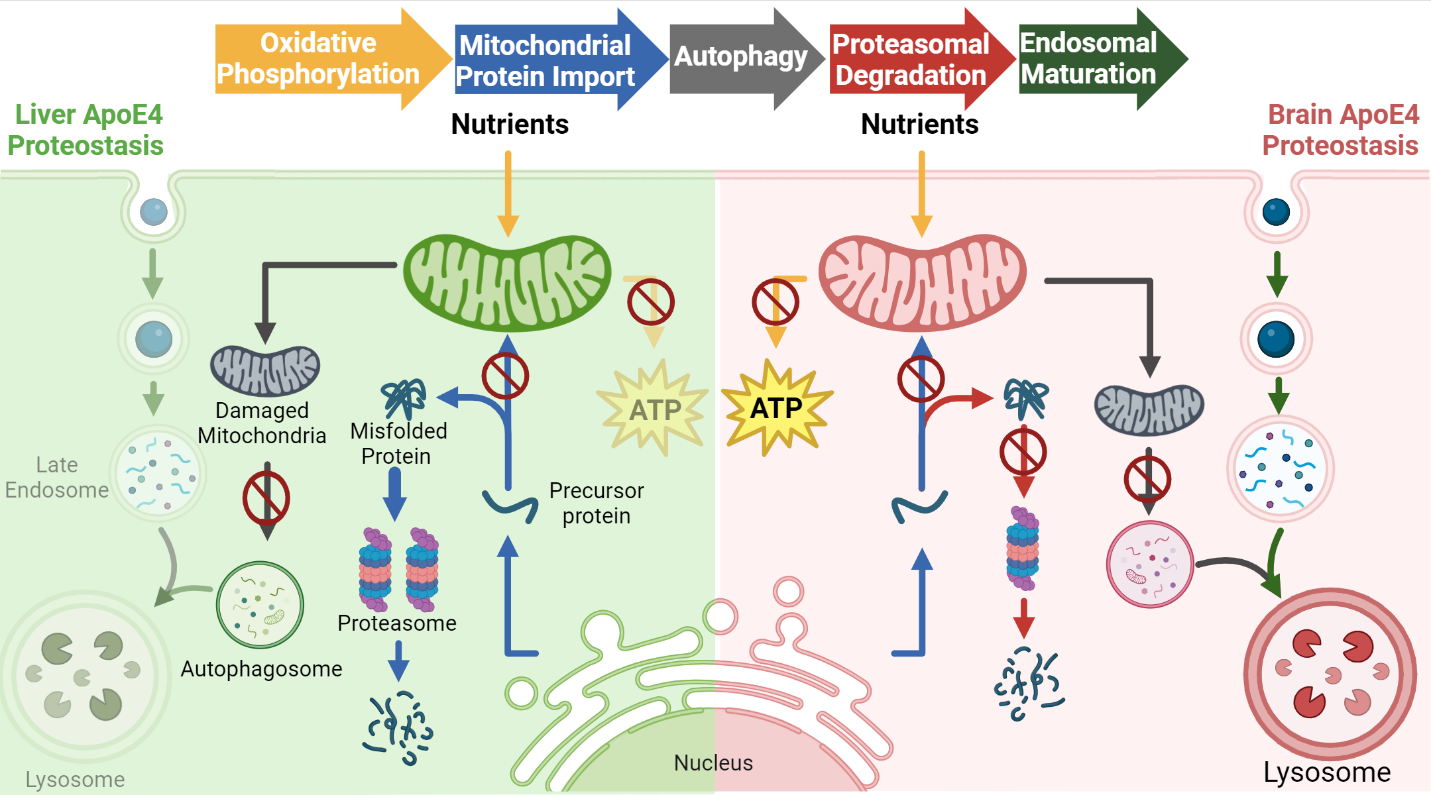

Supplementary Figure S7: ApoE4 change in brain (red) versus liver (green). The arrows are color coded to represent the different pathways impacted ApoE4 when compared to ApoE3
